# Structural and functional analysis of *Pseudomonas aeruginosa* PelA provides insight into the modification of the Pel exopolysaccharide

**DOI:** 10.1101/2024.06.28.601253

**Authors:** Jaime C. Van Loon, François Le Mauff, Mario A. Vargas, Stephanie Gilbert, Roland Pfoh, Zachary A. Morrison, Erum Razvi, Mark Nitz, Donald C. Sheppard, P. Lynne Howell

## Abstract

A major biofilm matrix determinant of *Pseudomonas aeruginosa* is the partially deacetylated α-1,4 linked *N*-acetylgalactosamine polymer, Pel. After synthesis and transport of the GalNAc polysaccharide across the inner membrane, PelA partially deacetylates and hydrolyzes Pel before its export out of the cell *via* PelB. While the Pel modification and export proteins are known to interact in the periplasm, it is unclear how the interaction of PelA and PelB coordinates these processes. To determine how PelA modifies the polymer, we determined its structure to 2.1 Å and found a unique arrangement of four distinct domains. We have shown previously that the hydrolase domain exhibits endo-α-1,4-*N*-acetylgalactosaminidase activity. Characterization of the deacetylase domain revealed that PelA is the founding member of a new carbohydrate esterase family, CE#. Further, we found that the PelAB interaction enhances the deacetylation of *N*-acetylgalactosamine oligosaccharides. Using the PelA structure in conjunction with AlphaFold2 modelling of the PelAB complex, we propose a model wherein PelB guides Pel to the deacetylase domain of PelA and subsequently to the porin domain of PelB for export. Perturbation or loss of the PelAB interaction would result in less efficient deacetylation and potentially result in increased Pel hydrolysis. In PelA homologues across many phyla, the predicted structure and active sites are conserved, suggesting that there is a common modification mechanism in Gram-negative bacterial species that contain a functional *pel* operon.

## INTRODUCTION

Biofilms are communities of microbial cells surrounded by an extracellular matrix composed of polysaccharides, proteins, surfactants, lipids, extracellular DNA (eDNA), and other nucleic acids (1, 2). The extracellular matrix is an important virulence factor that enables the embedded microbes to evade the host immune system and affects the efficacy of some antibiotics (3). The Gram-negative bacterium *Pseudomonas aeruginosa* predominantly exists as a biofilm and is commonly found in chronic infections, such as in the lungs of individuals with cystic fibrosis or burn wounds (4, 5). *P. aeruginosa* has the genetic capacity to produce three different exopolysaccharides, each of which is important for biofilm formation: alginate, Psl, and Pel (6). Pel provides structural support by forming interactions with eDNA in the biofilm matrix (7), aids in maintaining cell-to-cell interactions (8), interacts with abundant anionic host polymers in the sputum of individuals with Cystic Fibrosis (9), protects cells against aminoglycoside antibiotics (8, 10), helps the biofilm-embedded cells remain associated with the surface under starvation conditions (11), and provides a competitive advantage in pellicle biofilms under oxygen-limited conditions (12).

Pel is an α-1,4 linked linear polysaccharide that exists in both cell-associated and cell-free (also referred to as matrix-associated) forms (13). While the structure of the cell-associated form has yet to be determined, the cell-free form of Pel is composed predominantly of dimeric repeats of galactosamine (GalN) and *N*-acetylgalactosamine (GalNAc) (14). The biosynthesis of Pel in *P. aeruginosa* occurs via a synthase-dependent pathway and requires the proteins encoded by the *pelABCDEFG* operon (15, 16). In the cytoplasm, the PelDEFG complex synthesizes and transports the growing polysaccharide across the inner membrane (17). Once in the periplasm, the large 948-residue protein PelA modifies Pel using its two enzymatic domains. PelA’s deacetylase domain selectively removes acetyl groups from the polysaccharide, rendering Pel cationic (18), while its glycoside hydrolase family 166 (GH166) domain hydrolyzes Pel. The hydrolase activity is required for the generation of the cell-free form of the polysaccharide, which is important for the biomechanical properties of the biofilm and *P. aeruginosa* virulence (13, 19). PelB is a large protein that is proposed to help guide mature Pel through the periplasm using its tetratricopeptide repeats (TPRs). The β-barrel domain of PelB exports Pel to the extracellular space with the aid of the oligomeric lipoprotein PelC, which is proposed to act as an electronegative funnel to further guide Pel to the outer membrane (20, 21).

Interactions between modification enzymes and the periplasmic TPRs of outer membrane porin proteins are a common feature of synthase-dependent exopolysaccharide biosynthetic pathways. For example, the modification and export of poly-β(1,6)-*N*-acetylglucosamine (PNAG) in *E. coli* and alginate in *P. aeruginosa* is coupled through the PgaB-PgaA and AlgX-AlgK-AlgE interactions, respectively (22–26). For the modification and export of Pel in *P. aeruginosa*, an interaction between PelA and TPRs nine through fourteen of PelB results in an increase and decrease in the activities of PelA’s deacetylase and hydrolase domains, respectively (21). However, how the PelA-PelB interaction coordinates the modification and export of Pel for use in the *P. aeruginosa* biofilm is not known. Additionally, without structural information, it remains unclear how the multi-domain, dual-active protein PelA binds to and modifies Pel before export.

Herein, we present the crystal structure of PelA from *Pseudomonas thermotolerans* (*Pt*PelA) and show that the protein has a unique four-domain structure. Using pseudo-substrates and α-1,4-GalNAc oligosaccharides, we demonstrate that *Pt*PelA and its orthologue *P. aeruginosa* PelA (*Pa*PelA) exhibit hydrolase and deacetylase activity and that the deacetylase activity is enhanced when *Pa*PelA is incubated with its interaction partner, *P. aeruginosa* PelB (*Pa*PelB). In conjunction with Alphafold2 (AF2) modelling, we propose a model wherein *Pa*PelB guides Pel to the deacetylase domain of *Pa*PelA prior to export for use in the *P. aeruginosa* biofilm, ultimately blocking hydrolysis. If the interaction with *Pa*PelB is perturbed or lost, we hypothesize that *Pa*PelA can then hydrolyze the polysaccharide. The high degree of structural conservation of PelA across Gram-negative species suggests that the hydrolase and deacetylase activities of its homologues are essential for bacterial species that contain a functional *pel* operon.

## RESULTS

### PelA has a compact four-domain structure

To gain insight into the structure and function of PelA, we undertook structural studies of *Pa*PelA and a thermotolerant orthologue from *P. thermotolerans* (72% amino acid sequence identity and 83% similarity to *Pa*PelA). While *Pa*PelA did not readily crystalize, a construct of *Pt*PelA excluding the predicted signal sequence (SS), encompassing residues 37-937, crystallized in various conditions. Diffraction data were collected for selenomethionyl-incorporated (SeMet) *Pt*PelA at the National Synchrotron Light Source (NSLS)-II to 2.1 Å, and the structure was solved using the single-wavelength anomalous diffraction (SAD) method (27). Iterative refinement resulted in a R_work_ and R_free_ of 18.17% and 22.64%, respectively (Table 1). Structural analysis of *Pt*PelA reveals four distinct domains (Fig. 1AB) that align well with the AF2 model of *Pa*PelA (*Pa*PelA^AF2^; Fig. S1; 1.6 Å root mean square deviation (RMSD) over 816 α-carbons) (28). As the quality of the electron density prevented us from building residues 83-105, 198, 321-323, 389-395, 417-420, 448, 588-614, and 703-718, both the *Pt*PelA and *Pa*PelA^AF2^ models were used during our structural analyses.

**Figure 1.**
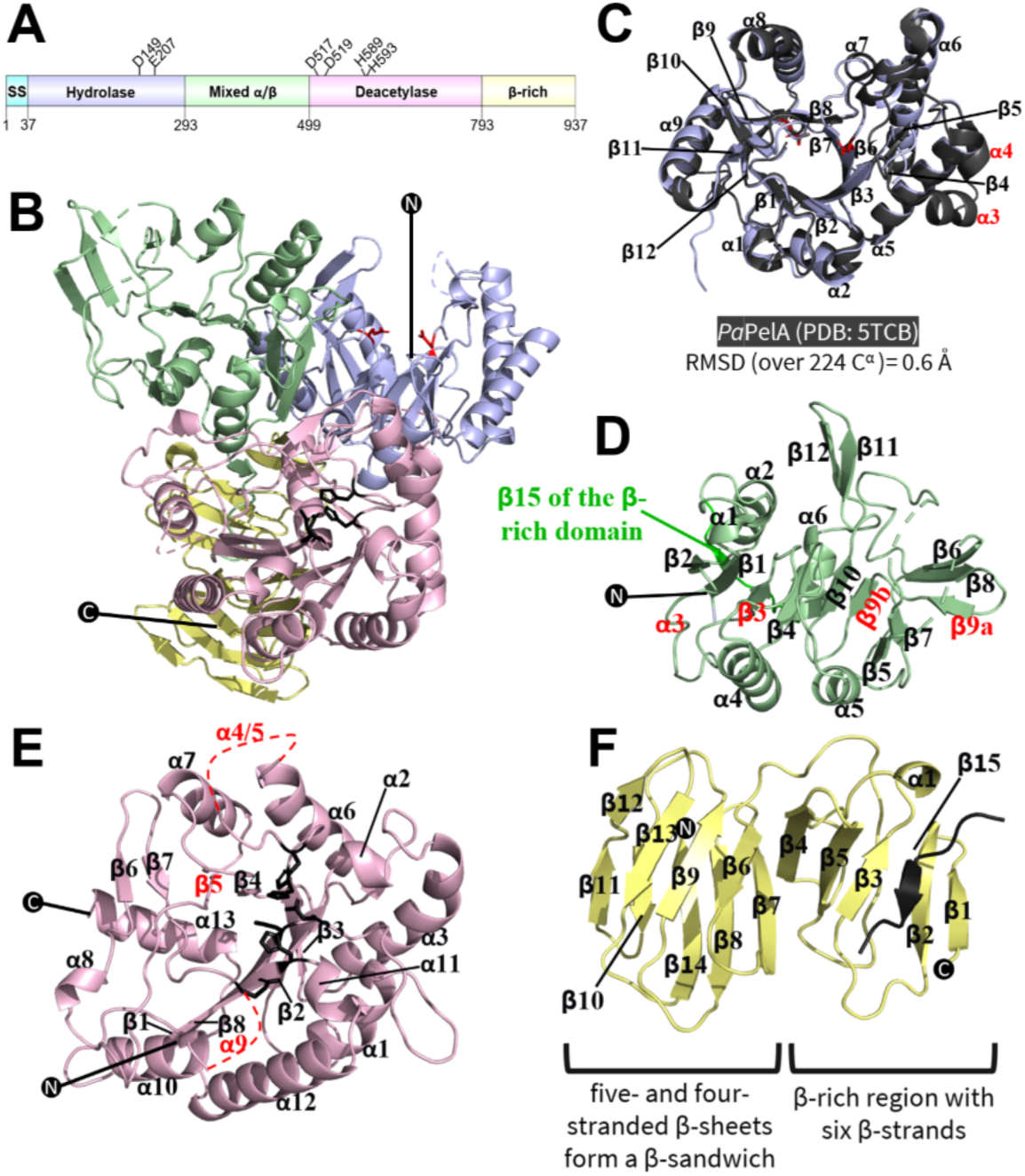
The full-length structure of *Pt*PelA. **(A)** Linear representation of *Pt*PelA domains based on the structure and available literature. The domain architecture was visualized using DOG 2.0 Illustrator of Protein Domain Structures (75). Amino acids that are required for enzymatic activity are highlighted. SS, signal sequence. **(B)** Cartoon representation of the crystal structure of residues 37-937 of *Pt*PelA shown in two opposing views. The domains as coloured as depicted in panel A. Amino acids required for the deacetylase and hydrolase activities are shown in black and red, respectively. N and C represent the N- and C-termini, respectively. The dashed lines represent regions of the protein that could not be built due to the poor quality of the electron density. **(C)** Comparison of the hydrolase domain of PelA as determined from the structure from *P. thermotolerans* (blue) and the structure of *Pa*PelA_H_ (5TCB, grey) with the secondary structure labelled. Secondary structural elements in the AF2 model of *Pa*PelA that are missing in the structure of *Pt*PelA are labeled in red. **(D)** The mixed α/β domain of *Pt*PelA. **(E)** The deacetylase domain of *Pt*PelA. **(F)** The β-rich domain of *Pt*PelA.

**Table 1.**
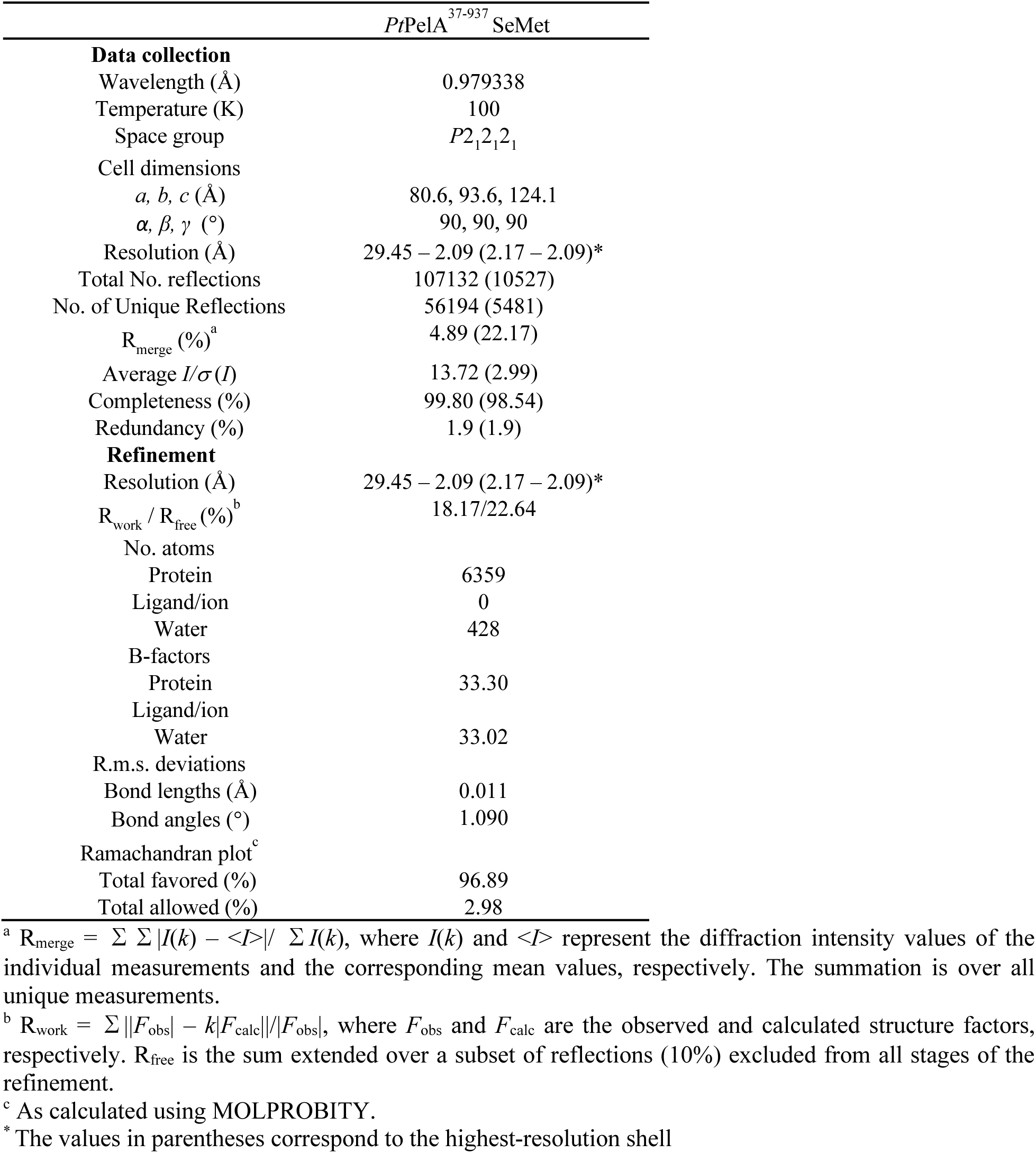
X-ray data collection and refinement statistics.

The N-terminal hydrolase domain, residues 37-292, has a β_12_/α_9_-barrel-fold that aligns closely to the structure of the isolated *Pa*PelA hydrolase domain (*Pa*PelA_H_, PDB 5TCB; 0.6 Å RMSD over 224 α-carbons) (Fig. 1C) (19). The hydrolase domains of *Pt*PelA and *Pa*PelA_H_ have a deep binding groove (Fig. 1B), which we have previously shown hydrolyzes GalNAc polymers preferentially but can also tolerate the presence of some GalN (14, 19).

Previous bioinformatics analyses have suggested that residues 303-409 in *Pa*PelA and residues 293-398 in *Pt*PelA are structurally similar to reductase enzymes (18). Upon analysis of the *Pt*PelA structure, we found that residues 293-498 form a mixed α/β domain with 12 β-strands (Fig. 1BD). Six β-strands (β2-β1-β3-β4-β10-β9b) run roughly parallel to each other through the center of the domain surrounded by five α-helices. In *Pa*PelA, the ninth β-strand forms the central strand of a five stranded β-sheet (β5-β7-β9a-β8-β6). In contrast, in the *Pt*PelA crystal structure, there is a discontinuity in β9 (Fig. 1D). While β9b is situated with β1-4 and β10, β9a is grouped with β5-8. There is a pair of β-strands (β11-12) that are surface exposed and physically adjacent to the deacetylase domain and an additional β-strand (denoted β15) that comes after the α6 helix, which contributes to the structure of the β-rich domain (Fig. 1DF). To provide insight into the function of this protein, we compared the structure of the mixed α/β domain of *Pt*PelA to the other experimentally determined structures using the DALI server (29). The top DALI hits reveal structural similarity to domains in a putative thua-like protein (PDB 4JQS; 5.1 Å RMSD over 168 α-carbons), the 2-amino-2-deoxyisochorismate synthase PhzE (PDB 3R75; 4.6 Å RMSD over 152 α-carbons) and an intraflagellar transport protein component IFT52 (PDB 5FMR; 5.9 Å RMSD over 120 α-carbons) (Fig. S2A) (29–32). These DALI hits have less than 15% sequence identity to *Pa*PelA and have several differences in their secondary structures, including a rearrangement of β5-8 and increased length of α1-3, a loss of β11-12 and a twisted conformation of β9-10, and changes to β6-9 and additional α-helices respectively (Fig. S2A). Examination of the function of these proteins suggests that this domain of PelA could bind carbohydrate and/or be important for mediating the interaction between PelA and PelB, as thua-like proteins are involved in the utilization of trehalose (30), while the region of PhzE and IFT52 that aligns to *Pt*PelA binds chorismite (32) and helps bridge interactions between the IFT-B1 and B2 components of this intracellular transport complex, respectively (31).

The deacetylase domain of *Pa*PelA (residues 520-804) has previously been predicted to have structural similarity to CE4 enzymes (18, 33). Amino acids 499-792 of *Pt*PelA reveal similarity to CE4 and CE18 enzymes (Table S1) (34, 35). The deacetylase domain of *Pa*PelA has the core (β/α)_7_-fold, which is characteristic of CE enzymes (Fig. 1E) (22, 34–37). In addition to the core (β/α)_7_-fold, there are a number of additional structural features. There are short α-helices between the second and seventh β/α pairs, a short β-strand and α-helix between the sixth β/α pair, and a 31-amino acid loop containing two small α-helices between the third β/α pair. Lastly, there is an α-helix after the last β/α pair that extends towards the final domain of PelA (Fig. 1E). A comparison of *Pa*PelA^AF2^ to *Pt*PelA reveals that the missing loops that encompass residues 588-614 and 703-719 in the *Pt*PelA structure are important for forming a 23 Å-deep, 44 Å-long Pel binding groove, which corresponds to the length of a (GalNAc)_9_ polymer (Fig. 2A) (38).

**Figure 2.**
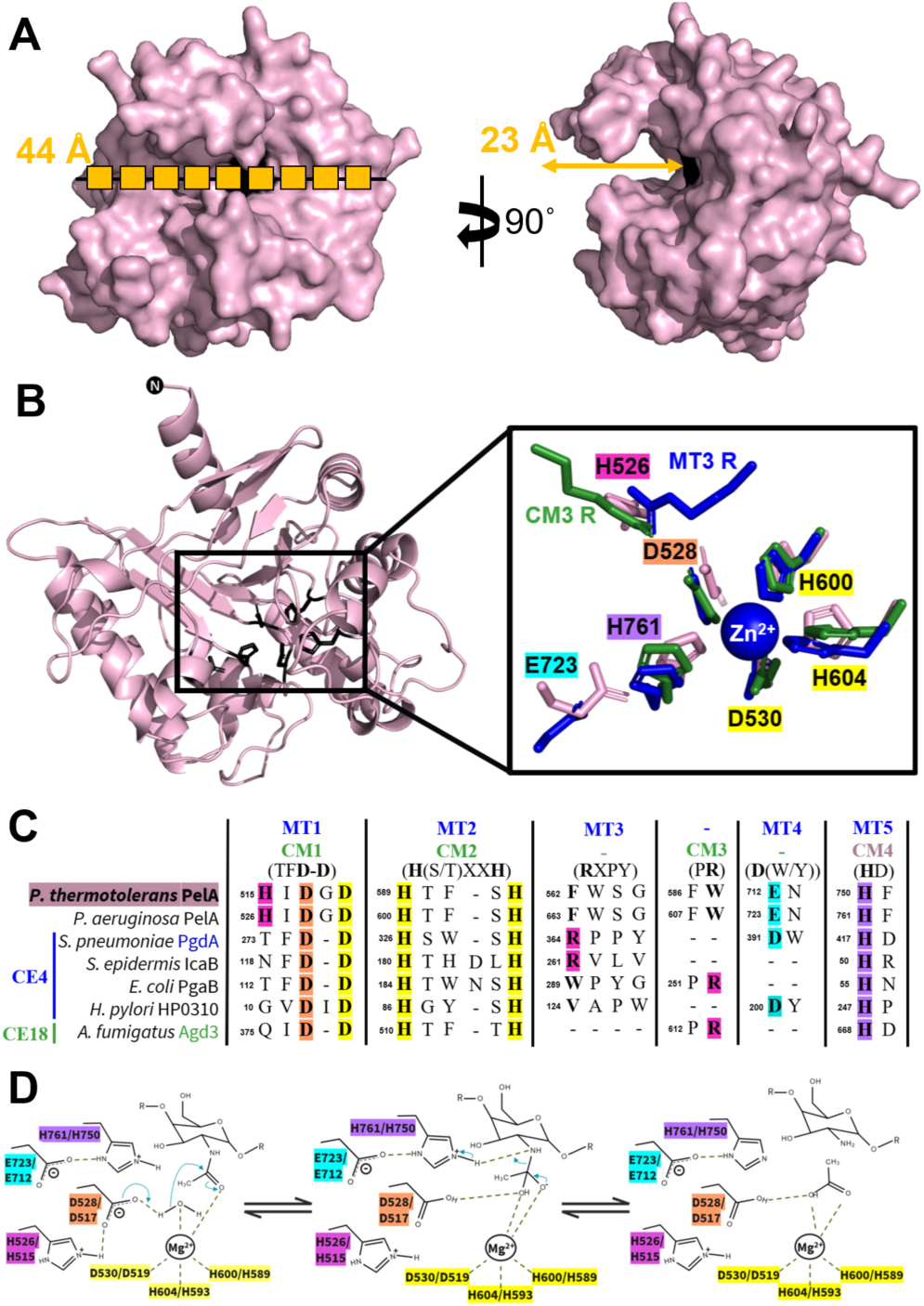
PelA is the founding member of a novel CE family. **(A)** Surface representation of the AF2 model of *Pa*PelA from two opposing views. The polymer in the first view of *Pa*PelA is representative of nine GalNAc residues (38). **(B)** The *Pa*PelA^AF2^ deacetylase domain (left). Inset of a structural alignment of the catalytic residues of *Pa*PelA (pink) with representative CE4 and CE18 enzymes: *S. pneumoniae* PgdA (2C1G, blue) and *A. fumigatus* Agd3 (6NWC, green), respectively. The zinc that co-crystallized with *Sp*PgdA is shown as a blue sphere. **(C)** Primary sequence alignment of the catalytic motifs MT1-5 and CM1-4 characteristic of CE4 and CE18 enzymes, respectively, as determined by structural alignment. The putative *Pa*PelA catalytic base (D528), metal coordinating triad (D530, H600, and H604), and putative catalytic acid (H761) are highlighted in orange, yellow, and purple, respectively. The aspartate that coordinates the catalytic acid (E723) and the histidine that is proposed to coordinate the catalytic base (H526) is shown in cyan and pink, respectively. **(D)** Proposed metal-dependent de-*N*-acetylation reaction for *Pa*PelA (first listed amino acid) and *Pt*PelA (second listed amino acid) based on the mechanisms for CE4 and CE18 enzymes and mutagenesis data.

*Pt*PelA also contains a 14-stranded β-rich region comprised of two distinct regions (residues 793-937) (Fig. 1F). The first β-rich region has six β-strands (β1-5 and β15), with a short α-helix positioned between β3 and β4. As described above, strand β15 is contributed from the mixed α/β domain (Fig. 1F). The second β-rich region consists of five- and four-stranded β-sheets that together form a β-sandwich (β6-14). The top DALI hits for the β-rich domain include the β-jelly domains of Xylanase B glycoside hydrolase (PDB 6KRN; 3.0 Å RMSD over 64 α-carbons), Dp0100 alginate lyase (PDB 6JP4; 5.6 Å RMSD over 120 α-carbons), and ChiD chitanase (PDB 5XSV; 2.7 Å RMSD over 64 α-carbons) (Fig. S2B) (39–42). While Dp0100 aligns to the entire β-rich domain of *Pt*PelA, Xylanase B and ChiD only align to the β-sandwich region of this domain. The nine-stranded β-jelly domain of Xylanase B has not been shown to bind xylan, but is required for the function of glycoside hydrolase family 30 enzymes (39, 40). Similarly, the 16-stranded β-sandwich domain of Dp0100 has not been shown to bind alginate (41). However, the presence of a β-jelly domain in alginate lyases is a common occurrence. The eight-stranded β-jelly domain of ChiD not only provides structural rigidity but this domain is also predicted to help generate a long chitin-binding site (42). Overall, we hypothesize that the β-rich domain of *Pt*PelA could provide structural stability to the protein and facilitate carbohydrate binding.

### PelA is the founding member of a new CE enzyme family

PelA deacetylase activity is essential for *P. aeruginosa* biofilm formation as chromosomal point mutation of residues in the putative deacetylase active site results in no adherence in a biofilm assay (18). To gain insight into PelA’s mechanism of action, we next compared the deacetylase domain to structures of previously determined carbohydrate esterase (CE) enzymes that are classified in the Carbohydrate Active enZyme (CAZy) database (18). While PelA was not currently part of any CE family, the DALI server revealed that PelA has some structural similarities to members of the CE4 and CE18 superfamilies (Table S1). For comparison purposes, we have used *Streptococcus pneumoniae* PgdA (*Sp*PgdA) and Agd3 as the representative CE4 and CE18 enzymes, respectively (34, 35).

While *Pa*PelA^AF2^ aligns poorly to *Sp*PgdA and Agd3 with RMSDs of 4.6 and 5.0 Å over 184 α-carbons, respectively, six significant structural differences between *Pa*PelA^AF2^ and *Sp*PgdA/Agd3 could be defined (Fig. 3). Four significant structural differences between *Pa*PelA^AF2^ and *Sp*PgdA contribute to a deeper and more elongated active site groove in *Pa*PelA^AF2^: an α/α-insertion preceding α6 (green), an extended loop containing α5 (orange), an extended loop preceding α1 (cyan), and a terminal α-insertion (dark grey) (Fig. 3A). Furthermore, *Pa*PelA has a β-insertion that follows β6, whereas there is a single terminal β-insertion in *Sp*PgdA (Fig. 3AB, brown). Similarly, an α/α-insertion preceding α6 (green), an extended loop containing α5 (orange), an extended loop preceding α1 (cyan), and a large, 64-amino acid loop insertion succeeding β2 (red) all contribute to an altered active site groove in *Pa*PelA compared to Agd3 (Fig. 3CD). The remaining differences between *Pa*PelA^AF2^ and *Sp*PgdA/Agd3 are in the active site, which will be outlined below.

**Figure 3.**
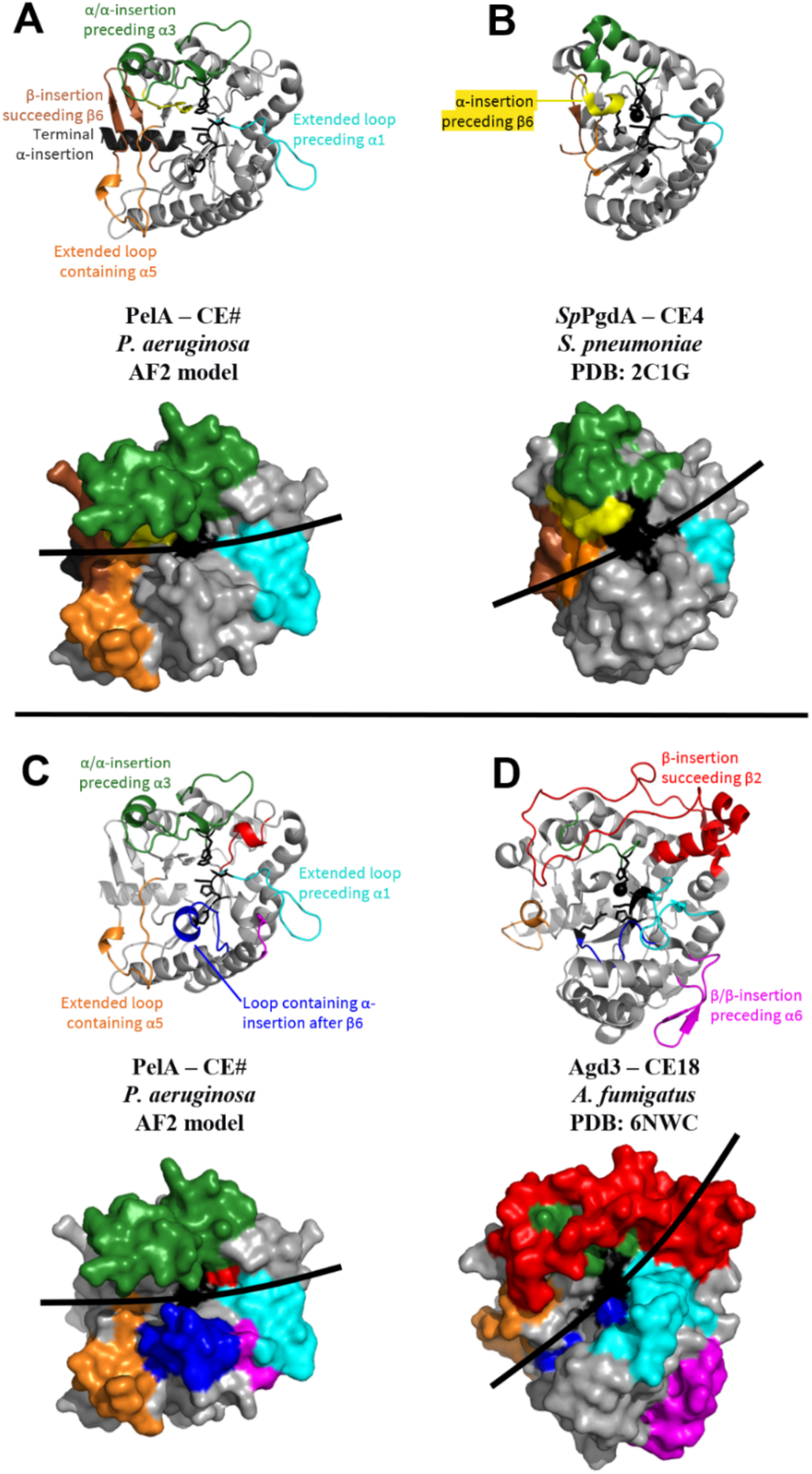
*Pt*PelAhas structural differences in comparison to CE4 and CE18 enzymes. **(A-B)** Cartoon and surface representations to compare the AF2 model of *Pa*PelA **(A)** and the structure of *Sp*PgdA **(B).** Regions where there are major structural differences are colored and labelled accordingly. Active site residues are shown in black. The black lines highlight the active site grooves. **(C-D)** Cartoon and surface representations to compare the AF2 model of *Pa*PelA **(C)** and the structure of Agd3 **(D).**

CE4 and CE18 family members have five and four canonical active site motifs, MT1-5 and CM1-4, respectively (Fig. 2BC). Residues D528 and D530 of *Pa*PelA are a part of MT1/CM1, while residues H600 and H604 are located in MT2/CM2. D528 is the putative catalytic base, whereas D530, H600, and H604 are predicted to be essential for coordinating the divalent cation required for deacetylation. The arginine residue found in MT3/CM3, which is responsible for activating the catalytic base in CE4 and C18 enzymes, is not conserved in *Pa*PelA. Compared to CE4 enzymes, *Pa*PelA is missing the α-insertion to orient MT3 in the active site (Fig. 3AB, yellow). The conserved arginine residue of Agd3 is situated in a loop that contains an α-insertion after β6 (Fig. 3CD, blue). While *Pa*PelA does have this insertion as well, the loop is oriented differently, and there is no conserved arginine residue in *Pa*PelA’s CM3. In contrast to CE4 and CE18 enzymes, *Pa*PelA contains a conserved histidine residue (H526), found in MT1, that aligns closely to the arginine residues in Agd3 and *Sp*PgdA (Fig. 2B). Finally, E723 in *Pa*PelA is a part of MT4, which is predicted to activate H761, the catalytic acid in MT5/CM4.

Due to the apparent lack of an activating arginine in the MT3/CM3 active site motif, *Pa*PelA likely operates *via* a modestly different mechanism than CE4 and CE18 enzymes. PelA potentially utilizes the histidine residue in its MT1/CM1 active site motif to activate the catalytic base.

### PtPelA is a metal-dependent CE enzyme with α-1,4-N-acetylgalactosaminidase activity

Previous studies of *Pa*PelA, its isolated hydrolase domain, and variants thereof have demonstrated that the enzyme exhibits esterase activity and is an α-1,4*-N-*acetylgalactosaminidase (18, 19, 43, 44). To confirm that *Pt*PelA is an active enzyme, we first probed for esterase activity using the pseudo-substrate acetoxymethyl-4-methylumbelliferone (AMMU) (44, 45). As anticipated, titration of AMMU into 1 µM of wild-type *Pt*PelA resulted in a concentration-dependent increase in esterase activity (Fig. S3). As CE enzymes require a divalent cation to stabilize the oxyanion tetrahedral intermediate during deacetylation (34, 35), we next sought to determine if *Pt*PelA’s esterase activity is dependent on the presence of metal and the identity of the metal. We first assessed the esterase activity of the wild-type protein and four alanine point mutants, including residues D517, D519, H589, and H593 (residues D528, D530, H600, and H604 in *Pa*PelA), which are implicated in the mechanism of deacetylation from our structural analyses. No detectable esterase activity was observed with any of the mutants (Fig. 4A), suggesting that a divalent metal is required and implicating D517 (residue D528 in *Pa*PelA) as the putative catalytic base. Differential scanning fluorimetry (DSF) revealed each of the mutants, with the exception of the H593A mutant, has comparable stability to the wild-type protein, suggesting that the changes observed in activity are not due to loss of stability (Table S2). Next, we pre-incubated the wild-type protein with metal chelators or the chloride salts of divalent cations and performed the AMMU esterase assay to determine the preferred metal (22, 34). While the addition of the dipicolinic acid (DPA) or ethylenediaminetetraacetic acid (EDTA) metal chelator resulted in a significant 2-fold reduction of esterase activity, *Pt*PelA still retained low levels of esterase activity (Fig. 4B). Exogenous addition of Mn^2+^ did not have a significant effect. While there was a trend towards increased activity when Ca^2+^ was added, this was not statistically significant. The addition of Mg^2+^ resulted in a significant, almost two-fold increase in esterase activity. Interestingly, the addition of Co^2+^, Cu^2+^, Ni^2+^, and Zn^2+^ resulted in a significant decrease in *Pt*PelA esterase activity (Fig. 4B). Collectively, these results show that PelA is a metal-dependent carbohydrate esterase.

**Figure 4.**
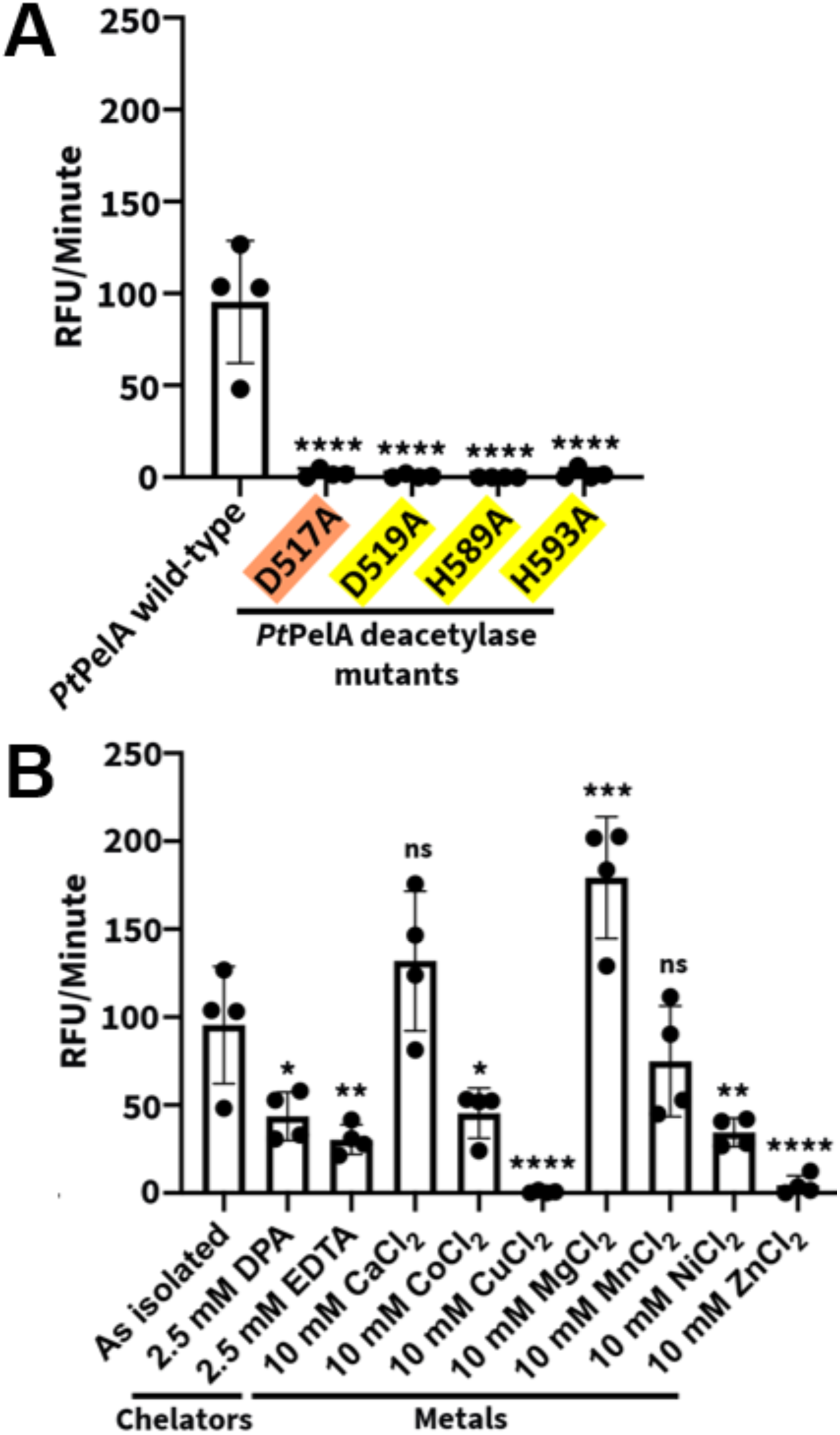
*Pt*PelA is a metal-dependent CE enzyme. **(A)** Detection of wild-type or mutant *Pt*PelA esterase activity using the AMMU pseudo-substrate. The putative catalytic base and metal coordinating triad are highlighted in orange and yellow, respectively. Statistical significance was calculated using an ordinary one-way analysis of variance with Dunnett’s multiple comparison test between wild-type and mutant *Pt*PelA. The error bars show the standard error of the mean for the four independent assays with two technical replicates. ****, p<0.0001; **, p<0.0021; *, p<0.0332; ns, not significant. **(B)** Detection of *Pt*PelA metal-dependent esterase activity using the AMMU substrate. Statistical significance was calculated using unpaired *t*-tests between *Pt*PelA as isolated and the other reaction conditions with chelator or metal chloride. AMMU, acetoxymethyl-4-methylumbelliferone.

Having confirmed that *Pt*PelA has esterase activity, we next sought to determine using a biofilm disruption assay whether the protein exhibited α-1,4-*N*-acetylgalactosaminidase activity (19, 43). Using *P. aeruginosa* PA14, which produces a Pel-dependent biofilm, we found that *Pt*PelA could disrupt preformed biofilms with a half-maximal effective concentration (EC_50_) of 78.5 ± 0.7 nM (Fig. 5AB). This is comparable to the EC_50_ for the recombinant hydrolase domain of *Pa*PelA (*Pa*PelA_H_) (88.3 ± 0.9 nM) (43). Alanine point mutants of residues required for *Pt*PelA hydrolase activity, D149 and E207, cannot disrupt the biofilms (Fig 5AB). To confirm these results, we assessed *Pt*PelA’s ability to hydrolyze a pool of α-1,4-GalNAc oligosaccharides. Digestion of this pool of oligosaccharides, which ranged from 5-mer (m/z = 1056.393 +/- 0.05) to 15-mer (m/z = 3087.110 +/- 0.10) (Fig 5C) (14, 19), was analyzed using matrix-assisted laser desorption/ionization-time of flight (MALDI-TOF) mass spectrometry (MS). Consistent with the results of the disruption assay, we observed a shift in the size range of the initial sample. After *Pt*PelA treatment, the oligosaccharides size ranged from 3-mer (m/z = 650.439 +/- 0.07) to 11-mer (m/z = 2274.803 +/- 0.10) (Fig. 5C and S4).

**Figure 5.**
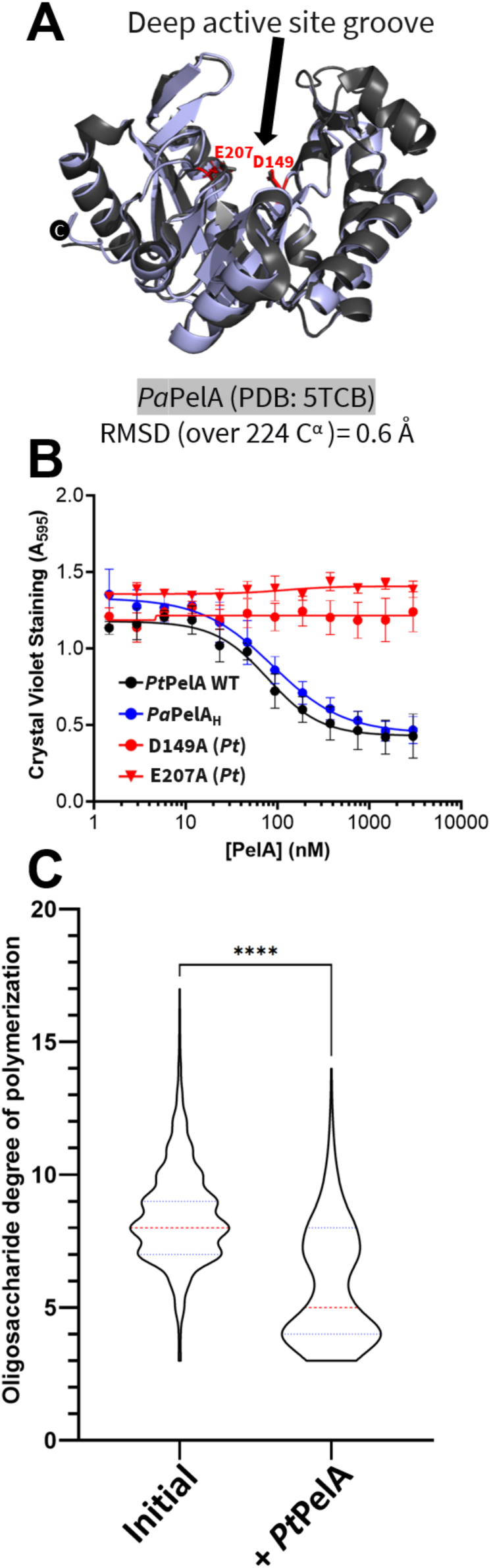
*Pt*PelA has α-1,4-*N*-acetylgalactosaminidase activity. **(A)** Structural comparison of the hydrolase domain of *Pt*PelA (blue) with the hydrolase domain of *Pa*PelA (5TCB, grey). Amino acids required for the hydrolase activity of *Pt*PelA (D149 and E207) and *Pa*PelA are shown in red and grey, respectively. **(B)** A biofilm disruption assay. Increasing concentrations of wild-type *Pt*PelA (black circle), *Pa*Pela_H_ (blue circle), or *Pt*PelA hydrolase domain mutants D149A (red circle) or E207A (red triangle) were added to pre-formed PA14 Pel biofilms. The error bars show the standard error of the mean for the average of three technical replicates in three independent assays. **(C)** Population statistical study of the MALDI-TOF MS enzyme spectra of the oligomers released due to hydrolysis after incubating wildtype *Pt*PelA. The data represent three biological replicates each with three technical replicates. Statistical significance was calculated using Dunnett’s multiple comparison test between the indicated reaction conditions. ****, p<0.0001. The median size of the oligosaccharides is shown by the dotted red line. The 25th and 75th quartiles are shown by the blue dotted lines. respectively.

Together, these results reveal that *Pt*PelA, like *Pa*PelA, exhibits both esterase and α-1,4-*N*-acetylgalactosaminidase activity. Given the low sequence identity to existing CE families, the deacetylase domain of *Pa*PelA and *Pt*PelA are the founding members of a new CE family, CE# (33).

### PaPelA CE function is regulated by PaPelB

We have previously shown that interaction with *Pa*PelB increases *Pa*PelA’s esterase activity in a *p*-nitrophenol assay and attenuates the hydrolysis of a preformed Pel-dependent *P. aeruginosa* biofilm (21). We hypothesized that the PelAB interaction resulted in less hydrolysis of Pel in the *P. aeruginosa* biofilm due to a potential increase in deacetylation. However, whether the presence and absence of PelB alters how PelA modifies Pel has not been characterized.

Given that *Pt*PelA has comparable esterase and hydrolase capabilities to *Pa*PelA (19, 43, 44), to be consistent with our previous studies, and given the availability of *Pa*PelA and *Pa*PelB constructs, our subsequent analysis focused on the *P. aeruginosa* enzymes. To examine both deacetylation and hydrolysis simultaneously and the effect PelB may have, a mix of purified α-1,4-GalNAc oligosaccharides of various lengths was added to 1 µM *Pa*PelA in the presence or absence of a soluble construct of *Pa*PelB that includes the TPRs and α-rich region that precedes the β-barrel porin domain (residues 47-880). The previously reported *Pa*PelA:*Pa*PelB ratio of 1:1.5 was used in the MALDI-TOF MS experiments (21). We monitored the potential hydrolysis and/or deacetylation reaction products over three time points. As the minimum substrate length for hydrolysis to occur is a heptamer (19), the relative ion proportion of the trimer to hexamer population was monitored and recorded as the reaction products. Only a minor amount of 5 and 6-mer was present in the initial sample (Fig. 6A). Incubation of the oligosaccharide mixture with *Pa*PelB did not alter the ion relative proportion of the end products (Fig. 6A and S5). In contrast, incubation with 1 µM of *Pa*PelA resulted in a constant increase in the trimer to hexamer product population (Fig. 6A). Addition of *Pa*PelB did not significantly change the population of hydrolysis products produced at any of the time points (Fig. 6A). We also tested the isolated hydrolase domain of *Pa*PelA (*Pa*PelA_H_) under similar conditions. Use of *Pa*PelA_H_ led to a similar increase in the 3- to 6-mer product population as observed with the full-length enzyme (Fig. 6B). Our results indicate that interaction with *Pa*PelB does not affect the hydrolase activity of *Pa*PelA or the isolated *Pa*PelA_H_ domain under the conditions tested.

**Figure 6.**
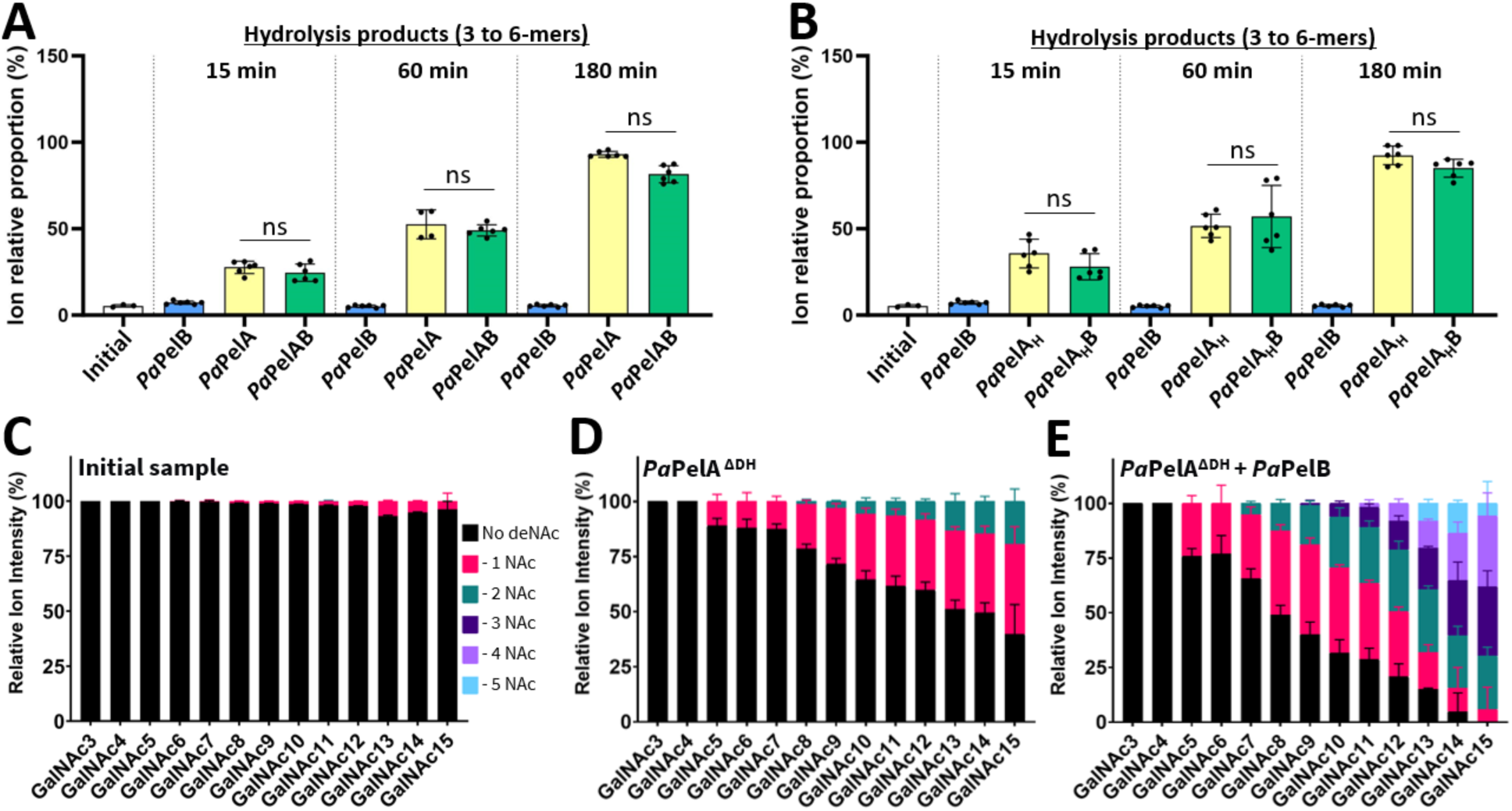
Interaction with *Pa*PelB does not affect *Pa*PelA hydrolase activity but increases the deacetylation of α-1,4-GalNAc oligosaccharides by *Pa*PelA. **(A-B)** The ion relative proportion of the MALDI-TOF MS enzyme spectra of the 3 to 6-mer oligomers released due to hydrolysis after incubating wildtype *Pa*PelA **(A)** or *Pa*PelA_H_ **(B)** in the absence or presence of *Pa*PelB. The data represent three biological replicates each with two technical replicates. Statistical significance was calculated using Kruskal Wallis multiple comparison tests between the indicated reaction conditions. Ns, not significant. **(C-E)** MALDI-TOF MS analysis of the deacetylation products for the oligosaccharides in isolation **(C)**, and after incubating *Pa*PelA^DH^ in the absence **(D)** or presence **(E)** of *Pa*PelB.

Under the conditions of the experiment, no oligosaccharide deacetylation was observed. The initial oligosaccharide sample contained a small (<3%) population of GalN residues (Fig S6A), and as anticipated this population didn’t change upon incubation with *Pa*PelB. However, in the presence of both *Pa*PelA and *Pa*PelAB, hydrolysis of the oligosaccharide leads to the loss of detectable GalN structures above the background noise of the spectra. As this data suggested that the deacetylase activity is slower than hydrolysis under the conditions tested, we generated a double hydrolase-inactivated mutant, *Pa*PelA^ΔDH^ (D160A E218A), to explore deacetylase activity. Using this enzyme, deacetylation events could be observed when 15 µM enzyme was incubated with the oligosaccharides for 24 hours. As *Pa*PelB is not stable when concentrated to 22.5 µM, we were unable to use the same 1:1.5 *Pa*PelA:*Pa*PelB ratio we used previously to examine the hydrolysis reaction and opted for a *Pa*PelA:*Pa*PelB ratio of 1.5:1. To determine the minimal length substrate required for deacetylation, the α-1,4-GalNAc oligosaccharides substrate mixture contained trimers and 15-mers. We found that the longer oligosaccharides are partially deacetylated (Fig. 6C), which is due to the nature of the purification and extraction process used to obtain the oligosaccharides. The shortest substrate that was affected by *Pa*PelA^ΔDH^ was a pentamer (Fig. 6D). The frequency and relative proportion of deacetylation events increased with oligosaccharide length. In contrast to the observed hydrolase activity, adding even a sub-stoichiometric amount of *Pa*PelB to *Pa*PelA^ΔDH^ enhanced the deacetylase activity (Fig. 6E). While the minimal substrate length required for deacetylation did not change, a marked increase in the relative proportion of oligosaccharide being deacetylated and the frequency of deacetylation events was observed (Fig. 6E). For example, in the absence of *Pa*PelB, only 60% of the 15-mer population was deacetylated with a maximum of two deacetylation events on the same oligomer. In the presence of *Pa*PelB, the whole population of 15-mer was deacetylated and up to 5 deacetylation events could be observed (Fig. 6E). Tandem mass spectrometry (MSMS) structural characterization of select deacetylated oligosaccharides was unable to reveal the exact position of the deacetylation events, highlighting the heterogeneity of the oligosaccharide processing by *Pa*PelA *in vitro* (Fig. S6BC). However, despite our inability to locate the first deacetylation event, it is important to note that the addition of *Pa*PelB did not alter the MSMS fragmentation trace obtained, demonstrating that the addition of *Pa*PelB did not modify how *Pa*PelA process the oligosaccharides (Fig. S6BC). Collectively, these results suggest that *Pa*PelB can directly affect the rate of deacetylase activity without changing the fundamental properties of minimal substrate length or sites of deacetylation.

### Full-length homologues of PelA are present in many Gram-negative bacterial phyla

When submitting the structure of *Pt*PelA to the DALI server, only structures of the isolated hydrolase or deacetylase domains of enzymes were identified (29). These results suggest that the domain organization in the full-length structure of *Pt*PelA is unique with respect to experimentally solved structures. As the AF2 database provides us with an opportunity to search for structural models of PelA homologues across all phyla, we next ran a non-redundant tertiary structure similarity search of *Pa*PelA^AF2^ using FoldSeek (Table S3.1) (46). We identified full-length homologues in 96 bacterial species that had >85% amino acid coverage and >45% sequence similarity as identified using ClustalW. We also searched a non-redundant database for homologues of *Pa*PelA using BLASTP (47), which searches based on sequence similarity. BLASTP identified 177 bacterial species with >85% amino acid coverage and >38% sequence similarity (Table S3.2). Aside from one Gram-positive species, all full-length homologues of *Pa*PelA identified in our FoldSeek and BLASTP searches were from Gram-negative species.

The sequences of *Pa*PelA and the homologues identified using FoldSeek were used to generate a maximum likelihood tree in Geneious, which identified 16 different bacterial phyla (Fig. 7A). One-third of hits were in the gammaproteobacterial phylum that *P. aeruginosa* and *P. thermotolerans* are a part of (Table S4). Betaproteobacterial species such as *Neisseriaceae*, *Rastonia*, and *Cupriavidus* contained about one-fourth of the full-length *Pa*PelA homologues. While four of the sequences came from unclassified phyla, most of the other sequences came from the aquificota, epsilonproteobacteria, and verrucomicrobiota phyla (Table S4). Interestingly, the homologues identified using BLASTP only identified eight unique bacterial phyla, with 33% and 63% of the hits corresponding to the betaproteobacterial and gammaproteobacterial phyla, respectively.

**Figure 7.**
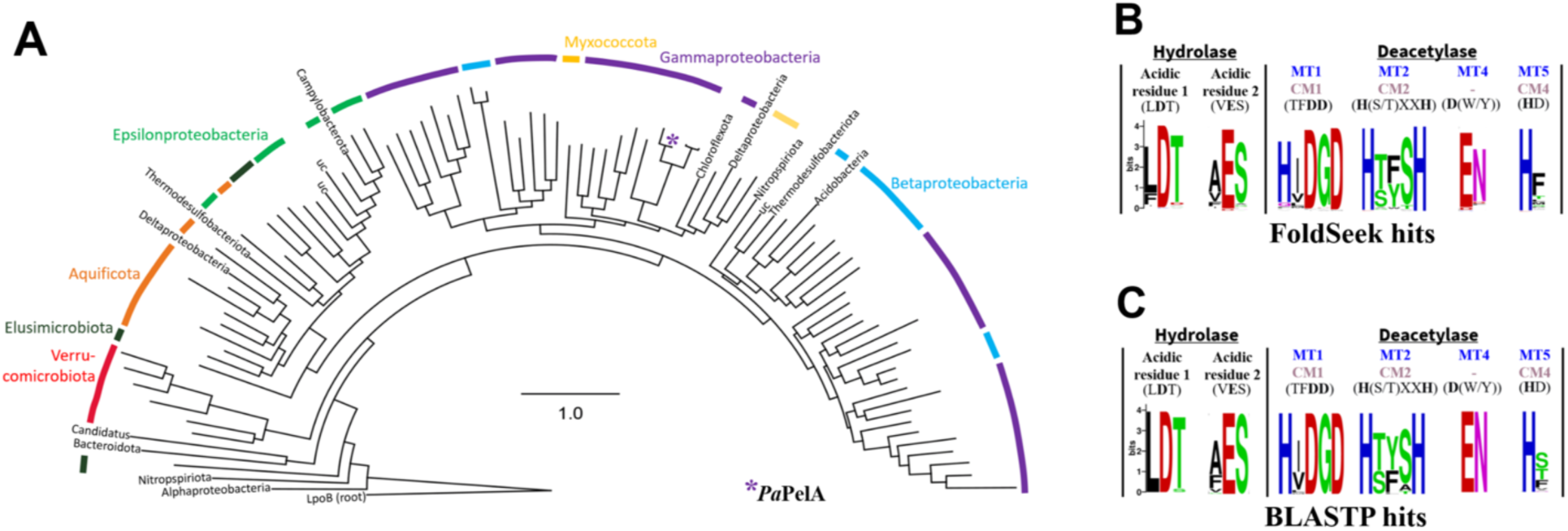
Full-length PelA is distributed across multiple Gram-negative phyla and has conserved active site residues. **(A)** Phylogenetic distribution of PelA among bacterial taxa. The circular tree represents the approximately-maximum-likelihood distance between 96 unique PelA sequences, which was constructed using FastTree and *Escherichia coli* LpoB as an outgroup. PelA sequences were identified by running a FoldSeek search of *Pa*PelA. **(B-C)** Multiple sequence alignment of *Pa*PelA homologues identified through FoldSeek **(B)** or BLASTP **(C)** and visualized using WebLogo 3. The catalytic motifs MT1-5 and CM1-4 are characteristic of CE4 and CE18 enzymes.

Using BioCyc, we found that only 21% (20/96) and 43% (76/177) of the *Pa*PelA homologues identified using FoldSeek and BLASTP, respectively, were linked to a *pelABCDEFG* operon. The FoldSeek and BLASTP data resulted in the identification of eight and seven new species, respectively, with *pel* or *pel*-like operons that had not been identified in our previous phylogenetic analyses (Table S5) (48). Considering that 5% and 44% of the *Pa*PelA homologues identified using FoldSeek and BLASTP were in unclassified species for which full genomic information is unavailable, more of these homologues may be in *pel* or *pel*-like operons. While the genomic context for many of the homologues identified in our searches is not available, every PelA homologue that we could search the genome for is in a *pel* operon. This strongly suggests all of the PelA homologues are probably associated with *pel* operons. Alignment of *Pa*PelA to its full-length homologues revealed conservation of the hydrolase and deacetylase active site amino acids with only minor differences occurring in the logos obtained from our FoldSeek and BLASTP data sets (Fig. 7BC). Due to the high degree of structural conservation, these results suggest that the hydrolase and deacetylase activities of the full-length PelA homologues could be essential in bacterial species that contain the *pel* operon. Additional work will be needed to determine if the *pel* operons are functional and if the PelA-PelB interaction remains relevant in these less-studied organisms.

### AF2 PaPelAB complex predicts PelA’s deacetylase and mixed α/β domains interact with PelB

While we know that the PelA-PelB interaction alters how PelA modifies Pel, it remains unclear which domain(s) of *Pa*PelA is/are essential for the interaction. Although there is a crystal structure of TPRs 8-11 of *Pa*PelB (21), the full-length structure of PelB has not been experimentally determined. To provide insight into how *Pa*PelA interacts with *Pa*PelB, we generated AF2 models of *Pa*PelB and the *Pa*PelAB complex (Fig. S7A). Examination of the AF2 *Pa*PelB structure and bioinformatics analyses suggest that amino acids 1-36 form an inner membrane α-helix rather than the hypothesized signal sequence (Fig. S7B) (21). As anticipated, amino acids 69-726 and 876-1193 encompass TPRs 1-19 and the β-barrel porin domain, respectively (Fig. 8A). While amino acids 37-68 and 727-910 were previously predicted to be unstructured regions (21), the AF2 model predicts with low confidence a single α-helix and an α-rich region for these residues, respectively (Fig. 8A and S7B). While the ten α-helices in the α-rich region do not form canonical ∼34-amino acid long TPR pairs (49), they do supercoil and form an electronegative portal that leads to the β-barrel porin domain (Fig. 8B).

**Figure 8.**
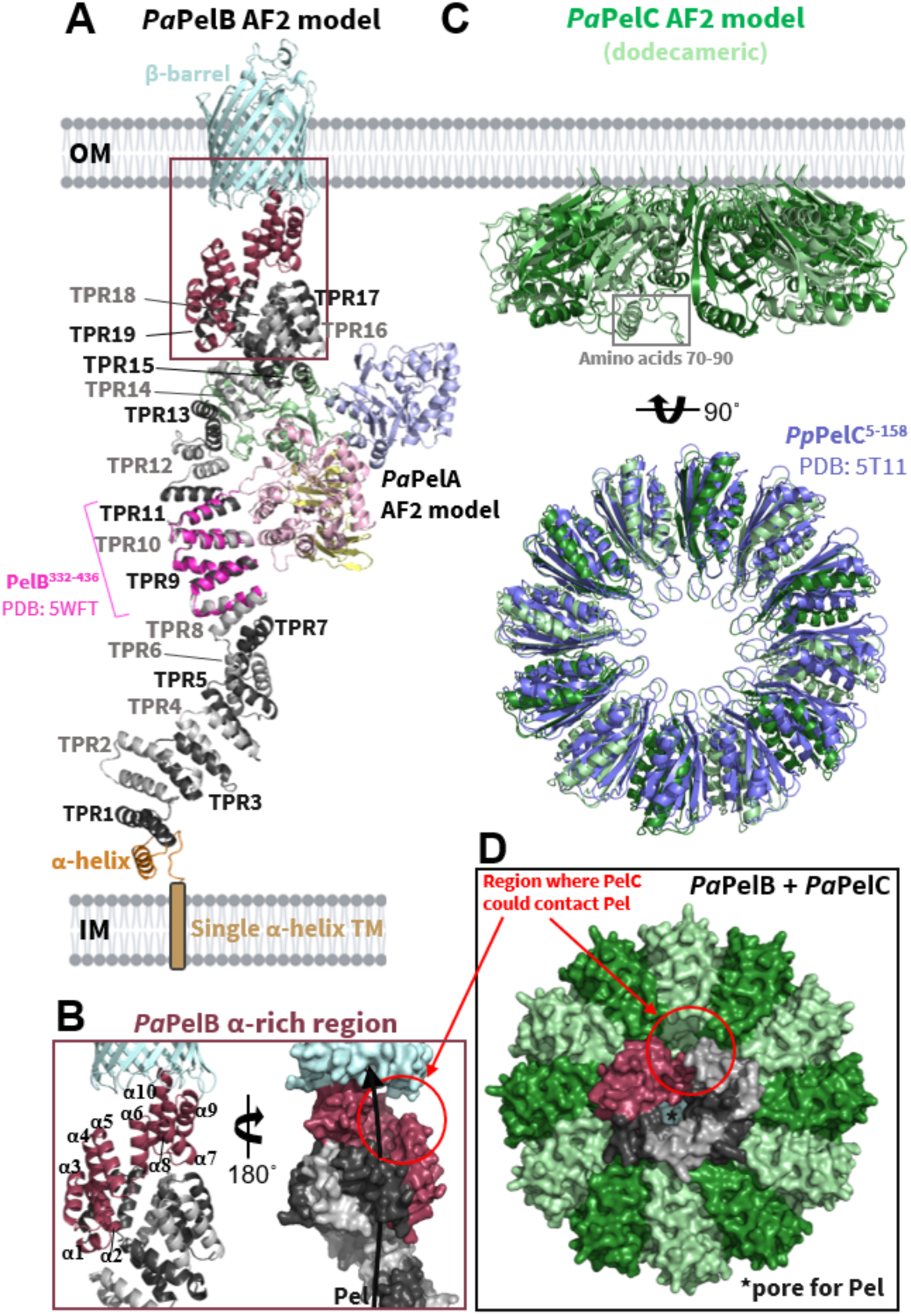
Structural models of *Pa*PelB, *Pa*PelC, and the *Pa*PelBC complex. **(A)** AF2 model of full-length *Pa*PelB complexed with *Pa*PelA (colored by domains as in Fig. 1). *Pa*PelB is colored by domain and aligned to the structure of *Pa*PelB^332-436^ (5WFT, magenta). **(B)** The α-rich region of *Pa*PelB from two opposing views. The black arrow indicates the proposed trajectory of Pel through PelB. **(C)** Alignment of the dodecameric AF2 prediction of *Pa*PelC with dodecameric *Pp*PelC (blue; RSMD of 1.5 Å over 136 Cα for the monomers). **(D)** The AF2 model prediction of amino acids 600-1193 of *Pa*PelB (colored as in panel A) complexed with the 12-subunit ring of *Pa*PelC (light and forest green). IM, Inner membrane, OM, outer membrane.

In keeping with our previous data that had shown that TPRs 9-14 of *Pa*PelB are required for the outer membrane localization of *Pa*PelA (21), the AF2 model of the PelAB complex predicts an interaction between the deacetylase domain of *Pa*PelA and TPRs 9-15 of *Pa*PelB (Fig. 8A) (21). The mixed α/β domain of *Pa*PelA is also predicted to interact with TPRs 13-19 of *Pa*PelB. This suggests that the function of the mixed α/β domain of PelA is likely to help coordinate *Pa*PelAB complex formation. In the AF2 *Pa*PelAB model, the hydrolase and β-rich domains of *Pa*PelA do not interact with *Pa*PelB (Fig. 8A).

### PaPelB and PaPelC interact to form the outer membrane Pel biosynthetic machinery

Based on structural studies, we have previously hypothesized that the dodecameric lipoprotein PelC interacts with PelB and acts as an electrostatic funnel to attract cationic Pel and guide it toward the porin domain for export (20). Given the AF2 models of *Pa*PelB and the *Pa*PelAB complex, we hypothesized that PelC may also interact with PelA and, therefore, could potentially influence its activity. Attempts to predict the entire *Pa*PelAB-dodecameric PelC complex (>4200 residues) were unsuccessful, and therefore, we modelled amino acids 600-1193 of *Pa*PelB with dodecameric *Pa*PelC using AF2 (Fig. 8CD and S7C). To note, Alphafold3 produced the same models, with the exception of a kink after the eighth TPR of *Pa*PelB (50). We found that the AF2 dodecameric *Pa*PelC model aligns closely with our structure of *Paraburkholderia phytofirmans* PelC (*Pp*PelC) (RSMD of 1.5 Å over 136 Cα for the monomers) and that the AF2 model conserved the protomer-protomer interface found in the crystal structure (Fig. 8C) (20). Minor differences arise as a flexible loop, and an α-helix that is periplasmically facing could not be built in the *Pp*PelC structure due to the poor quality of the electron density in this region (amino acids 70-90 in *Pa*PelA; grey box in Fig. 8C). The AF2 complex prediction suggests that the electropositive exterior surface of TPRs 17-19 and the α-rich region of *Pa*PelB mediate the interaction with the electronegative interior pore of *Pa*PelC (Fig. S8). While we have previously hypothesized that the negative pore of PelC is essential for guiding positively charged, deacetylated Pel to the OM β-barrel domain of PelB, the *Pa*PelBC complex suggests that there is only a small region of PelC that could encounter Pel (red circle in Fig. 8D). As there is a small gap in the α-rich region of *Pa*PelB that could expose Pel to the periplasm, the wider electronegative pore of *Pa*PelC closes over this region and would result in a fully enclosed, electronegative pore for Pel to travel to the outer membrane porin domain of *Pa*PelB (red circles in Fig. 8CD and green circles in Fig. S8). Consistent with our previous study showing that the residues that contribute to the electronegativity of *Pa*PelC are essential for Pel-dependent *P. aeruginosa* biofilm formation, the AF2 complex prediction suggests that the electronegative pores of both *Pa*PelB and *Pa*PelC are required for the export of deacetylated Pel.

### Model showing the projected path of Pel through the PaPelABC complex

To gain insight into how Pel is modified and exported, we next built a composite model of the *Pa*PelABC complex by combining our PelAB and PelBC models (Fig. 9A). This model shows that PelA and PelC do not interact and that PelC is, therefore, unlikely to affect PelA function. Our *Pa*PelABC model suggests that after synthesis and translocation of the polymer across the cytoplasmic inner membrane by PelDEFG, Pel interacts with TPRs 1-9 of PelB, an interaction which helps guide the polymer through the periplasmic space and delivers it to PelA. The *Pa*PelA-*Pa*PelB interaction orients the deacetylase domain so that it can readily bind to the fully acetylated GalNAc polysaccharide. The interaction of PelB and PelA *via* TPR 9-15 exposes an electronegative groove that we propose guides the now positively charged, partially deacetylated product towards the outer membrane for export. If the polymer follows this path, it will not be exposed to the hydrolase domain (Fig. 9BC).

**Figure 9.**
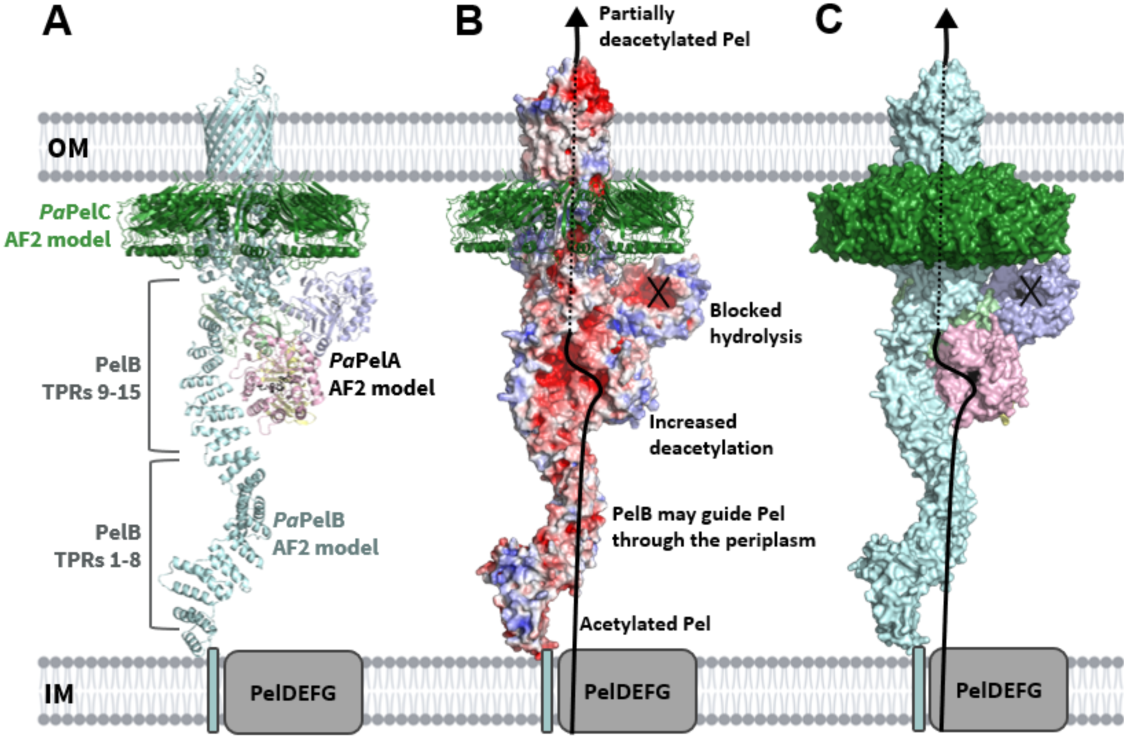
Model of the *Pa*PelABC outer membrane Pel modification and secretion complex. **(A)** AF2 model of full-length *Pa*PelB (light cyan) complexed with *Pa*PelA (colored by domains as in Fig. 1). AF2 model of amino acids 600-1193 of *Pa*PelB complexed with the 12-subunit ring of *Pa*PelC (green) were aligned to the *Pa*PelAB complex prediction. The fragment of *Pa*PelB used for the *Pa*PelBC prediction was then excluded from this figure. **(B)** Cartoon representation of dodecameric *Pa*PelC and the surface representation of *Pa*PelAB from panel A as colored by electrostatics, which were calculated by APBS in PyMol and visualized from -5 (blue) to 5 (red) kT/e. The solid black line indicates the proposed trajectory of Pel export while *Pa*PelA and *Pa*PelB interact. The dotted black line indicates when the polymer is enclosed by the pore formed by the *Pa*PelAB complex. **(C)** Surface representation of panel A. The active site amino acids of the deacetylase and hydrolase domains of *Pa*PelA are shown in black.

## DISCUSSION

In this study, we present the structural characterization of the Pel polysaccharide modification enzyme, PelA. We determined the crystal structure of PelA from *P. thermotolerans* to 2.1 Å, revealing a unique four-domain architecture (Fig. 1). Bioinformatics analyses of *Pseudomonas* PelA homologues reveal conservation of the catalytic residues and predicted structures (Table S3; Fig. 7), suggesting that PelA modifies a GalNAc-rich polysaccharide in bacterial species that contain a *pel* operon. Using sequence analysis and enzymatic assays, we found that *Pt*PelA and its homologues define a new class of metal-dependent carbohydrate esterases, CE#, with distant homology to the CE4 and CE18 families (Fig. 2, 3 and 4). Importantly, we show using MALDI-TOF MS that the interaction of *Pa*PelA with *Pa*PelB results in increased deacetylation of α-1,4-GalNAc oligosaccharides (Fig. 6CDE) but does not directly affect the proportion of hydrolytic products (Fig. 6AB).

Deacetylation of exopolysaccharides is an important modification in Gram-negative and -positive bacterial biofilms (22, 35–37). *Pa*PelA deacetylase activity is essential for biofilm formation, as chromosomal mutation of deacetylase active site residues in *P. aeruginosa* PA14 results in impaired biofilm formation (18). We found that *Pt*PelA’s esterase activity was enhanced with the addition of magnesium ions (Fig. 4B). While metal dependency studies of *Pa*PelA were not possible due to the poor stability of the protein, intracellular concentrations of magnesium in *P. aeruginosa* are typically high (100-500 mM) suggesting that *Pa*PelA’s activity may also be enhanced by magnesium (51–53). However, CE enzymes are often promiscuous in their metal dependency (22, 33–35, 52, 53). As many CE4 enzymes are zinc-dependent and/or have been co-crystallized with zinc, other metals, such as zinc, could enhance the deacetylase activity of *Pa*PelA *in vivo* (22, 34–36, 53, 54). Furthermore, as *P. aeruginosa* and *P. thermotolerans* reside in different environmental niches, it is possible that the metal utilized by *Pa*PelA may vary from the results seen for *Pt*PelA (55, 56).

*Pa*PelA lacks the mostly conserved arginine in the MT3 and CM3 active site motifs of CE4 and CE18 enzymes, respectively (Fig. 2BC). In CE4 and CE18 family members, this arginine is proposed to activate the aspartate that functions as the catalytic base. There is some variation across CE4 enzymes regarding both the location of this arginine in the primary sequence and the orientation it adopts in the active site. For example, compared to *Sp*PgdA, the conserved arginine in *E. coli* PgaB is positioned in the CM3 motif that is characteristic of CE18 enzymes and is oriented differently in the active site with respect to the catalytic base from MT1 (22). Enzymes that are classified as CE4 family members but lack any apparent activating arginine residue are not active, such as PgdA from *H. pylori* (*Hp*PgdA) (57). *Hp*PgdA does not have an arginine residue in its active site and, under the conditions tested, did not show activity on the *N*-acetylated polysaccharides that are typically used by peptidoglycan deacetylases (35, 62, 63). While *Pt*PelA and *Pa*PelA lack the arginine that activates the catalytic base in CE4/CE18 enzymes, our analysis reveals that they are active deacetylases. Examination of the *Pt*PelA structure and bioinformatic analyses of the PelA homologues identified using FoldSeek and BLASTP suggest that the conserved histidine in the MT1/CM1 performs this function. With the exception of this difference in the residue that activates the catalytic base, *Pa*PelA has all of the CE4/CE18 consensus residues in its active site motifs. We propose a mechanism for *Pa*PelA and CE# enzymes wherein the water that is coordinated by a divalent cation is deprotonated by the catalytic base, D528 (Fig. 2D). The resulting nucleophile then attacks the acetate group of acetylated Pel, generating an oxyanion tetrahedral intermediate that is stabilized by the divalent cation. The catalytic acid, H750, then donates a proton to the intermediate, resulting in the release of acetate from Pel (Fig. 2D).

While there were many Gram-negative hits identified in our list of PelA homologues, *Streptococcus dysgalactiae* was the only full-length, Gram-positive homologue (Table S4). Unfortunately, genomic data for this homologue is not available and so we can not determine whether the protein is situated in a *pel* operon. However, we know from our previous phylogenetic studies that *pel* operons are widespread across Gram-positive species (58) and in these operons, PelA is typically split into two proteins: a deacetylase with a mixed α/β and β-jelly roll domain, and a separate glycoside hydrolase. Both of these proteins play a role in Pel-dependent *B. cereus* biofilm formation. *B. cereus* PelA is essential for biofilm formation, while deletion of the hydrolase gene results in increased biofilm biomass are (58), a phenotype also observed in *P. aeruginosa* when hydrolase activity is abrogated (13). We hypothesize that changes to the domain architecture and structure of Gram-positive PelA homologues may stem from the lack of an outer membrane and loss of *pelB* and *pelC* from the operon.

Our previous studies showed that full-length *Pa*PelA, but not the isolated hydrolase domain, interacts with *Pa*PelB (21). The AF2 prediction of the *Pa*PelAB complex supports these data, as only the deacetylase and mixed α/β domains of *Pa*PelA are involved in the interaction interface (Fig. 9). As AF2 is unable to predict conformational changes, the current AF2 model only enables a mechanism for the deacetylation of Pel to be predicted. In *E. coli*, the dual-active protein PgaB first partially deacetylates the PNAG polymer (26). Deacetylated PNAG is then predicted to wrap around PgaB until it encounters the hydrolase domain before being exported by the OM protein PgaA (26). It is, therefore, possible that a shift in the mixed α/β and/or deacetylase domains of *Pa*PelA could alter the binding groove of the protein such that hydrolysis can occur. While it is unclear whether the structure of *Pa*PelB is rigid, it is also possible that *Pa*PelB can undergo conformational changes in response to periplasmic stress or from an unknown interaction partner, which could ultimately affect how the protein interacts with *Pa*PelA. Consequently, our previous studies and the *in vitro* functional studies presented herein of the *Pa*PelAB complex may not account for the conditions that would favor a different conformational state. As the hydrolase domain is important for generating the cell-free form of Pel and thus controls biofilm biomechanics and plays a role in modulating *P. aeruginosa* virulence, future work is required to understand if there are conditions under which hydrolysis is activated. Furthermore, determining the structure of the *Pa*PelAB complex in the presence and absence of Pel will be required to understand how *Pa*PelA can deacetylate and hydrolyze the polymer before secretion from the cell.

The *Pa*PelAB interaction resulted in increased deacetylation of α-1,4-GalNAc oligosaccharides (Fig. 6CDE). It is important to keep in mind that hydrolysis of α-1,4-GalNAc oligosaccharides dominates over deacetylation *in vitro*, and that the *Pa*PelAB complex may not bind the full-length Pel polymer in the same way during biosynthesis. Despite our previously published findings that the addition of *Pa*PelB increased the hydrolytic activity of *Pa*PelA in a preformed biofilm disruption assay, *Pa*PelB did not affect the hydrolysis of α-1,4-GalNAc oligosaccharides (Fig. 6AB). While the substrate for *Pa*PelA in the preformed *P. aeruginosa* biofilm contained partially deacetylated Pel, fully acetylated oligosaccharides were used as the substrate in the MALDI-TOF experiments herein. Therefore, it is likely that the acetylation state of Pel affects how the PelAB interaction regulates hydrolysis. This data supports our previous study showing that *Pa*PelA_H_ hydrolyzes acetylated Pel more easily than mature, partially deacetylated Pel (14). Alternatively, it is possible that additional interaction partners or different conditions would be required to elicit a change in the hydrolysis of Pel by *Pa*PelA and its interaction with *Pa*PelB.

Despite being less efficient than the hydrolase domain, the deacetylase domain of *Pa*PelA likely acts on the Pel polysaccharide first since Pel is likely guided directly to the deacetylase domain from the PelDEFG inner membrane complex. Additionally, deacetylation is prioritized as it is required to export Pel through the electronegative exit channel formed by *Pa*PelB and *Pa*PelC (Fig. S8) (18, 20, 21). Disruption of the interaction between *Pa*PelA and *Pa*PelB, release of PelA, or an alteration in how these proteins interact, could lead to less efficient deacetylation of the polymer and an increase in the percentage of GalNAc content in Pel. As the hydrolase domain of *Pa*PelA is more efficient at cleaving acetylated Pel (14, 19), PelA could then rapidly hydrolyze the polysaccharide (13). Determining how Pel binds to the deacetylase and hydrolase domain active sites of *Pa*PelA and the mechanisms of deacetylation and hydrolysis of the polysaccharide with respect to the *Pa*PelA-*Pa*PelB protein complex will be crucial for understanding how the Pel polysaccharide is modified for use in the biofilm matrix of *P. aeruginosa* and other Pel-producing bacteria.

## METHODS

### Bacterial strains, plasmids, and growth conditions

A complete list of bacterial strains and plasmids used in this study can be found in Table S6. Terrific broth (TB) was used for the growth of the *E. coli* strains. TB contained, per litre of ultrapure water, 47.6 g of terrific broth and 0.4 % (w/v) glycerol. No-salt lysogeny broth (NS-LB) was used for the growth of the *P. aeruginosa* strains. NS-LB contained, per liter of ultrapure water, 10 g of tryptone and 5 g of yeast extract. Semisolid media was prepared by adding 1.5% (w/v) agar to LB. Where appropriate, kanamycin (Kan) at 50 µg/mL was added to growth media for *E. coli*.

### Standard molecular methods

All basic microbiological and molecular biological techniques were performed using standard protocols. Genomic DNA was isolated using BioRad InstaGene Matrix. Plasmid isolation and DNA gel extraction was performed using purification kits purchased from BioBasic. Restriction enzymes, DNA ligase, alkaline phosphatase, and DNA polymerase were purchased from Thermo Scientific. Primers used in this study were obtained from Sigma-Aldrich (Table S7). Site-directed mutagenesis of plasmids was carried out using the Agilent QuikChange Lightning site-directed mutagenesis kit. Transformation of *E. coli* was performed using standard protocols. Sanger sequencing to confirm the sequence of plasmids and chromosomal mutations was performed at The Center for Applied Genomics (TCAG), Toronto.

### Cloning of Recombinant proteins

The nucleotide sequence of *P. thermotolerans* H165_RS0111390 PelA (*Pt*PelA; accession WP_026146567) was obtained from The *Pseudomonas* DataBase and used to order a gene synthesis product cloned into the pET24a vector (BioBasic) (59). The PRED-TAT server indicated that *Pt*PelA is processed via the general secretion system (Sec) and has a signal sequence from residues 1-22. To obtain a soluble construct, the *Pt*PelA pET24a construct was used as a template to amplify the DNA of *Pt*PelA without the predicted signal sequence and at the start of the hydrolase domain (amino acids 37-937) (60). The restriction enzymes NcoI and XhoI were introduced and the gene was ligated into the pET28a expression vector encoding an N-terminal hexahistidine (His_6_) tag. This generated the soluble *Pt*PelA construct used in the studies herein.

The site-directed mutants of *Pt*PelA and *Pa*PelA without its predicted signal sequence (amino acids 47-948 for *Pa*PelA; accession Q9HZE4) were generated using the QuikChange Lightning site-directed mutagenesis kit (Stratagene). The sequence of all vectors was confirmed using DNA sequencing.

### Expression and purification

Expression and purification of wild-type *Pa*PelA (amino acids 47-948), the hydrolase construct of *Pa*PelA (amino acids 47-303), and *Pa*PelB (amino acids 47-880) has been described previously (18, 21, 43). Expression and purification of the double hydrolase mutant of *Pa*PelA was carried out as described for the wildtype protein. The expression of N-terminally His_6_ tagged *Pt*PelA was achieved through the transformation of an expression vector into *E. coli* BL21 CodonPlus (DE3) cells, which was then grown in 1 L TB containing 50 µg/mL Kan at 37 °C. The cells were grown to an OD_600_ of 0.8-1.0, at which point isopropyl-β-D-1-thiogalactopyranoside (IPTG) was added to a final concentration of 1 mM to induce expression. The induced cells were incubated for 20 hr at 18°C before being harvested via centrifugation at 6000 x *g* for 25 min at 4 °C. The pellet was stored at -20 °C until used.

The cell pellet was thawed and resuspended in 40 mL of wash buffer (20 mM HEPES pH 8, 300 mM NaCl, 10% (v/v) glycerol, 1 mM tris(2-carboxyethyl)phosphine (TCEP), and 20 mM imidazole) containing one SIGMAFAST protease inhibitor ethylenediaminetetraacetic (EDTA)-free mixture tablet (Sigma). The resuspended cells were lysed with the sonicator 3000 (Misonix) for a processing time of 5 min. The pulse was applied on for 5 s and off for 10 s. The output level was 6.5. The insoluble lysate was removed by centrifugation at 25000 x *g* for 30 min at 4 °C. The supernatant was loaded onto a 5 mL nickel-affinity column preequilibrated with 50 mL of wash buffer. The beads were then washed with 150 mL of wash buffer. Bound His_6_-tagged protein was eluted using 50 mL of wash buffer containing 250 mM imidazole. Eluted protein was concentrated to 2 mL by centrifugation using Amicon Ultra centrifugal filters (Millipore) with a 50-kDa molecular weight cut-off (MWCO) at 3500 x *g* at a temperature of 4 °C. The protein was further purified using size exclusion chromatography with a HiLoad 16/60 Superdex 200 gel filtration column (GE Healthcare) into a final buffer of 20 mM HEPES pH 8, 150 mM NaCl, 10 mM TCEP, and 5% (v/v) glycerol. After gel filtration, protein fractions corresponding to greater than 95% purity on a 12% sodium dodecyl sulphate-polyacrylamide gel electrophoresis (SDS-PAGE) gel were pooled and concentrated using a 50-KDa MWCO Amicon Ultra centrifuge filter. Protein samples were divided into aliquots and stored at -80 °C until use. The mutant constructs were expressed and purified as per the wild-type protein, and their folding and stability were assessed using DSF (Table S2).

Expression of the selenomethionyl-incorporated (SeMet) *Pt*PelA in a minimal medium was performed with a previously described protocol using B834 Met^-^ *E. coli* cells (Novagen) and subsequently purified using the protocol for the wild-type protein (61).

### Crystallization, data collection, structure solution and refinement

Purified *Pt*PelA was screened using the sitting-drop vapor diffusion method with the Crystal Gryphon Robot (Art Robbins Instruments). Commercial sparse-matrix crystal screens MCSG1-4 (Microlytic) and JCSG TOP96 (Rigaku) were used to screen crystallization conditions. Trials were set up at room temperature and at a protein concentration of 1.5, 3 or 5 mg/mL. In the Intelli 96-3 Shallow Well plates (Hampton), 1 µL drops at a 1:1 protein to mother liquor ratio were set up over a reservoir containing 100 µL of the crystallization solution. Native crystals were visible in many conditions after 5 days. Crystals formed in 12.5% (w/v) PEG3000 and 0.1 M sodium citrate pH 5.7 (TOP96 condition #C4) were harvested, cryoprotected with 20% (v/v) glycerol, and vitrified in liquid nitrogen. 2.7 Å resolution data were collected at the NSLS-II using the AMX beamline (17ID-1) beamline at −173 °C.

SeMet *Pt*PelA was screened for crystal hitsas previously described, except for increasing the drop size from 1 to 2 µL. Crystals were visible in many conditions after 5 days. Crystals that formed in 0.17 M ammonium sulfate, 0.085 M sodium citrate pH 5.6, 25.5% (w/v) PEG4000, and 15% (v/v) glycerol (TOP96 condition #A10) were harvested and vitrified in liquid nitrogen. Se-SAD data were collected at the NSLS-II using the AMX beamline (17ID-1) at −173 °C (0.2° oscillations, 360°, 0.01 s/image). Datasets were merged using XDS (62). The structure was solved using the SHELX suite (63). A total of twelve out of sixteen selenium sites in *Pt*PelA had significant anomalous signal and were located using SHELXE (63). The polyalanine trace was initially able to build 381 residues, and the preliminary phased electron density map was used for manual model building in COOT (64). Residues 750-791 were first built into the density. The AF2 model of *Pt*PelA along with the remaining alanine trace were used to build into the density. There was no interpretable density for the His_6_ tag and for the following residues: 83-105, 198, 321-323, 389-395, 417-420, 448, 588-614, and 703-718. Iterative refinement of the model was performed in real space using Coot and through PHENIX.REFINE (65, 66).

### Structural analysis tools

Individual protein and complex structural model predictions were generated using the AF2 Protein Structure Database. All structural figures were generated using PyMOL (The PyMOL Molecular Graphics System, Version 1.2, Schr_Ö_dinger, LLC) and ChimeraX 1.7 (Resource for Biocomputing Visualization, and Informatics RBVI, UCSF) (67). The DALI server was used to compare the structures of full-length and the individual domains of *Pt*PelA with existing structures in the Protein Data Base (PDB) (29). All structural biology applications used in this project were compiled and configured by SBGrid (68).

### Multiple sequence analysis

The amino acid FASTA sequences of the analyzed CE enzymes (PelA, *P. aeruginosa* PAO1; PgdA, *S. pneumoniae* – accession Q8DP63; HP0310, *Helicobacter pylori* ATCC 700392/26695 – accession O25080; IcaB, *Staphylococcus epidermidis* 35984 – accession Q5HKP8; PgaB, *E. coli* K12 – accession P75906; and Agd3, *Aspergillus fumigatus* Af293 – accession Q4WX15) were retrieved from the *Pseudomonas* Genome DataBank (Pseudomonas.com) or UniProt (69). Multiple sequence alignment was performed using the Clustal Omega server (70).

### AMMU esterase assay to detect metal dependency

This assay was performed as previously described (44). Thawed wild-type and mutant *Pt*PelA was diluted to 1 µM in the assay buffer (50 mM HEPES pH 7 and 75 mM NaCl) and kept on ice. The protein was then serially diluted from 1.11 µM to 1.1 nM. The AMMU substrate was dissolved in dimethyl sulfoxide (DMSO) prior to use. In PCR strip tubes, 20 µL of protein was incubated with 2.5 µL of MilliQ water, 100 mM metal, or 25 mM metal chelator for 10 min. Reactions were initiated when 2.5 µL of AMMU was added to the enzyme reactions solutions to a total reaction volume of 25 µL. From the PCR strip tubes, 10 µL from the reaction solution were added to a 384-well black bottom plate (Corning 3820) in duplicate and centrifuged at 3000 RPM for 30 s. Reaction progress was monitored in real time by measuring the RFU at 30 s intervals over 10 min at room temperature. The λ_emm_ and λ_ext_ used were 330 and 450 nm, respectively.

The background hydrolysis was monitored and subtracted from the enzyme-catalyzed reactions. All assays were performed in quadruplicate using a BioTek Synergy Neo2 plate reader (Agilent Technologies). Prism (GraphPad Software Inc.) was used for all analyses.

### Microtiter dish biofilm disruption assays

This assay was performed as previously described (43). An overnight culture of PA14 was grown in LB-NS and normalized to an OD_600_ of 0.5, then diluted to an OD_600_ of 0.005 in 1 mL of LB-NS. 100 µL of normalized cells were added to the wells of 96-well anionic Corning CellBind plates and incubated statically for 24 h at 25 °C. Following incubation, nonattached cells were removed, and the plate was rinsed thoroughly three times with water. 142.5 µL of phosphate-buffered solution (PBS) was added to the wells. The protein was serially diluted from 60 to 0.06 µM in PBS, and 7.5 µL of each protein sample was added to the wells (n=3 for each concentration). After incubating for 1 h at room temperature with shaking, the plate was rinsed thoroughly three times with water. The plates were then stained with 150 µL of 0.1% (w/v) crystal violet for 10 min at room temperature. The plate was rinsed, adhered crystal violet was solubilized in 150 µL of 100% (v/v) ethanol for 10 min at room temperature, and then the absorbance at 595 nm was measured to quantify adhered cells. Prism (GraphPad Software Inc.) was used for all analyses.

### Biochemical characterization of enzyme activity by MALDI-TOF MS

For all enzymes, a mix of α-(1,4)-GalNAc oligomers of various degrees of polymerization was used as substrate. Oligosaccharides were prepared as previously described (19). Briefly, 24 h-old biofilms of *A. fumigatus* Af293 were incubated with 10 nM Sph3 in 0.1X PBS for 1 h. Released oligosaccharides were then purified on C18 Sep-pak column.

The hydrolase activity of *Pt*PelA was observed after incubating the α-(1,4)-GalNAc oligosaccharides with 1 µM of enzyme in 1X PBS for 1 h at 37 °C. The hydrolase activity of *Pa*PelA was observed after incubating the α-(1,4)-GalNAc oligosaccharides with 1 µM of enzyme in 20 mM HEPES pH 8, 150 mM NaCl, and 5% (v/v) glycerol for 15 min, 1 h, and 3 h at room temperature in absence or presence of 1.5 µM *Pa*PelB. The deacetylase activity of *Pa*PelA was observed after incubating the α-(1,4)-GalNAc oligosaccharides with 15 µM of *Pa*PelA double hydrolase mutant (E218A and D160A) in the same buffer used for wild-type enzyme for 24 h at 37 °C with or without 10 µM *Pa*PelB.

After each digestion, oligosaccharides were purified from the enzymes and salt on Hypersep HyperCarb Sep-pak columns and eluted with 100% CAN. Eluates were dried down under airflow and reconstituted in 10 µL of 0.2% trifluoroacetic acid (TFA) before being spotted onto a MALDI plate in 2,5-dihydroxybenzoic acid (DHB) matrix as previously reported. Analysis of the oligosaccharide population by MALDI-TOF MS was acquired in positive reflectron mode, accumulating at least 10000 shots on Bruker UltrafleXtreme equipment. Prism (GraphPad Software Inc.) was used for the final analyses.

### Homologue identification and analysis

*Pa*PelA^AF2^ was submitted to FoldSeek with an E-value cut-off of 0.03 and five iterations, resulting in the identification of 96 full-length sequences (71). The sequences were aligned by Clustal Omega in Geneious Prime 2023 (72). The alignment was then analyzed using FastTree v2 implemented within Geneious Prime 2023 using the *E. coli* lipoprotein LpoB (GenBank accession number WP_000164439.1) as an outgroup (73). The amino acid sequence of PelA from *P. aeruginosa* was also submitted to BLASTP with a cut-off value of 1e-03 with a sequence cap of 1000 (47). Only 177/1000 of the sequences identified in the BLASTP search were full-length and used in this study. When possible, BioCyc was used to identify Pel-like operons in the genomes of PelA homologs identified in the FoldSeek and BLASTP searches (74).

## Supporting information

Tabel S3

## Acknowledgements

We thank Greg Wasney from the SickKids Structural & Biophysical Core (SBC) facility for using the equipment and having constructive discussions.

## Author Contributions

JCVL and PLH conceptualization; JCVL and PLH formal analysis; JCVL, FLM, MAV, SG, RP, ZM, and ER investigation; JCVL and PLH writing original draft; All authors reviewed and approved the manuscript; DCS, MN, and PLH resources; DCS, MN, and PLH supervision; DCS, MN, and PLH funding acquisition; PLH project administration.

