## Supplemental Information for "Structural and functional analysis of *Pseudomonas aeruginosa* PelA provides insight into the modification of the Pel exopolysaccharide"

**Running title:** Modification of the Pel exopolysaccharide by PelA

**Keywords**: *Pseudomonas aeruginosa*, biofilm, Pel polysaccharide, crystallography, structure-function.

**Table S1. The deacetylase domain of *Pt*PelA$^{\$}$ is structurally similar to carbohydrate esterases belonging to families CE4 and CE18.**

| PROTEIN NAME/FUNCTION (IF KNOWN) | Reference | CE type as defined by CAZy | Species | Cation co-crystallized | RMSD (Å) | PDB | # of aligned residues | # of residues in protein | % identity |
|---|---|---|---|---|---|---|---|---|---|
| **Polysaccharide deacetylase*** | - | - | *Burkholderia pseudomallei* | - | 2.9 | 3s6o-C | 193 | 306 | 11 |
| **Peptidoglycan GlcNAc deacetylase *Sp*PgdA** | (1) | CE4 | *Streptococcus pneumoniae* | $Zn^{2+}$ | 2.4 | 2c1g-A | 158 | 384 | 14 |
| **4-alpha-glucanotransferase** | (2) | - | *Thermococcus litoralis* | $Ca^{2+}$ | 3.3 | 1k1x-A | 200 | 636 | 8 |
| **Chitin deacetylase** | (3) | CE4 | *Bombyx mori* | $Zn^{2+}$ | 3.2 | 5zns-A | 199 | 381 | 11 |
| **Pseudoenzyme BA3943*** | - | CE4 | *Bacillus anthracis* | - | 2.5 | 7bkf-A | 156 | 281 | 14 |
| **Alpha amylase glucoside hydrolase (GH57)** | (4) | - | *Thermococcus kodakaraensis* | - | 3.2 | 3n8t-A | 202 | 550 | 9 |
| **Glycoside hydrolase/deacetylase BDI3119*** | - | - | *Parabacteroides distasonis* | - | 5.8 | 3lm3-A | 203 | 434 | 7 |
| **Chitin deacetylase DA1** | (5) | - | *Vibrio cholerae* | $Zn^{2+}$ | 3.7 | 4nyu-A | 183 | 406 | 7 |
| **Deacetylase Agd3** | (6) | CE18 | *Aspergillus fumigatus* | $Zn^{2+}$ | 3.8 | 6nwz-A | 212 | 665 | 10 |
| **Polysaccharide deacetylase WBMS*** | - | - | *Bordetella bronchiseptica* | $Zn^{2+}$ | 2.8 | 3hft-A | 158 | 242 | 9 |
| **Uncharacterized protein ATU2773*** | - | - | *Agrobacterium tumefaciens* | - | 2.8 | 2qv5-A | 159 | 246 | 11 |
| **Alpha-mannosidase Ams1** | (7) | - | *Schizosaccharomyces pombe* | $Zn^{2+}$ | 3.3 | 7dd9-A | 189 | 1129 | 6 |
| **Deacetylase BH1492*** | - | - | *Bacillus halodurans* | $Zn^{2+}$ | 3.0 | 2nly-A | 160 | 216 | 13 |
| **Hypothetical protein PHO986*** | - | - | *Pyrococcus horikoshii* | - | 3.3 | 1v6t-A | 165 | 249 | 10 |
| **Hypothetical hydrolase BT_2193*** | - | - | *Bacteroides thetaiotaomicron* | - | 3.6 | 3sgg-A | 183 | 512 | 8 |
| **Deacetylase Ba0331** | (8) | - | *Bacillus anthracis* | $Zn^{2+}$ | 3.2 | 6go1-A | 139 | 318 | 10 |
| **Maltose-forming amylase Py04_0872** | (9) | - | *Pyrococcus sp. ST04* | - | 3.7 | 4cmr-A | 190 | 597 | 10 |
| **Protein EF3048*** | - | - | *Enterococcus faecalis* | - | 3.3 | 2i5i-A | 169 | 261 | 10 |
| **Putative polysaccharide deacetylase Bd3279** | (10) | - | *Bdellovibrio bacteriovorus* | $Zn^{2+}$ | 3.3 | 5jp6-A | 156 | 339 | 13 |
| **Putative polysaccharide deacetylase BACOVA_03992*** | - | - | *Bacteroides ovatus* | - | 4.8 | 4dwe-A | 174 | 460 | 7 |

$^{\$}$*Pt*PelA has 937 residues, 294 of which comprise its deacetylase domain.

*To be published

**Table S2. Melting temperatures of *Pt*PelA point mutants as determined by differential scanning fluorimetry (DSF).**

| ***Pt*PelA enzyme** | **Domain of mutation** | **$T_m$ (˚C)** |
|---|---|---|
| WT | n/a | 53.06 ± 0.19 |
| D149A | Hydrolase | 52.23 ± 1.34 |
| E207A | Hydrolase | 57.02 ± 1.15 |
| D149A E207A (double hydrolase mutant, DH) | Hydrolase | 52.89 ± 0.24 |
| D517A | Deacetylase | 55.35 ± 1.04 |
| D519A | Deacetylase | 53.82 ± 0.71 |
| H589A | Deacetylase | 49.37 ± 0.42 |
| H593A | Deacetylase | 44.99 ± 0.33 |

**Table S3. List of full-length PelA homologues identified through a database of AF2 models used for comparative modeling (FoldSeek) and through sequence similarity (BLASTP).**
Please see file: SI Table 3.

**Table S4. Summary of the phyla distribution of full-length PelA homologues from Table S3.**

| ***Bacterial phyla*** | **FoldSeek hits** | **BLASTP hits** |
|---|---|---|
| *Acidobacteria* | 1 | 0 |
| *Alphaproteobacteria* | 1 | 0 |
| *Aquificota* | 9 | 0 |
| *Bacillota** | 0 | 1 |
| *Bacteroidota* | 1 | 1 |
| *Betaproteobacteria* | 21 | 59 |
| *Campylobacterota* | 1 | 0 |
| *Candidatus* | 1 | 0 |
| *Chloroflexota* | 1 | 1 |
| *Deltaproteobacteria* | 2 | 1 |
| *Elusimicrobiota* | 4 | 0 |
| *Epsilonproteobacteria* | 7 | 0 |
| *Gammaproteobacteria* | 31 | 111 |
| *Myxococcota* | 3 | 2 |
| *Nitrospiriota* | 2 | 0 |
| *Thermodesulfobacteriota* | 2 | 0 |
| *Undetermined* | 4 | 0 |
| *Verrucomicrobiota* | 6 | 0 |
| *Zetaproteobacteria* | 0 | 1 |
| ***Total*** | **97** | **177** |

*The only Gram-positive hit.

**Table S5. Summary of newly identified species that contain *pel* or *pel*-like operons as identified from BioCyc analysis of the full-length PelA homologues from Table S3.**

| ***Method of identification*** | **Species name** |
|---|---|
| ***FoldSeek hits*** | *Acidihalobacter ferrooxydans* |
| | *Nautilia profundicola* AmH |
| | *Nitrosospira* sp. NpAV |
| | *Pandoraea thiooxydans* |
| | *Persephonella marina* EX-H1 |
| | *Stigmatella aurantiaca* DW4/3-1 |
| | *Sulfurihydrogenibium* sp. YO3AOP1 |
| | *Thermovibrio ammonificans* HB-1 |
| ***BLASTP hits*** | *Atopomonas hussainii* |
| | *Halomonas hamiltonii* |
| | *Halomonas lutescens* |
| | *Nitrosomonas communis* |
| | *Pseudomonas agarici* |
| | *Pseudomonas mangrovi* |
| | *Simplicispira suum* |

**Table S6. Bacterial strains and plasmids used in this study.**

| Strain/Plasmid | Description | Source |
|---|---|---|
| ***E. coli* strains** | | |
| DH5α | Cloning strain; F⁻ Φ80*lacZ*ΔM15 Δ(*lacZYA-argF*) U169 *recA1 endA1 hsdR17* ($r_K^-$, $m_K^+$) *phoA supE44* λ⁻ *thi*-1 *gyrA96 relA1* | Invitrogen |
| BL21-CodonPlus | Protein expression strain; F⁻ , *ompT hsdS* (rB⁻ mB⁻) dcm⁺ Tet$^R$ galλ (DE3) endA [*argU proL* Cam$^R$] | Stratagene |
| B834 Met⁻ | SeMet protein expression strain: F⁻ *ompT hsdS*$_B$($r_B^-$ $m_B^-$) *gal dcm met* (DE3) | Novagen |
| ***P. aeruginosa* strains** | | |
| PA14 | Wild-type strain | M.R. Parsek |
| PAO1 Δ*wspF* Δ*psl* $P_{BAD}$*pel* | PAO1 Δ*wspF* (in-frame); Δ*pslBCD* (polar); *araC*-$P_{BAD}$ inserted upstream of *pelABCDEFG* | (11) |
| ***A. fumigatus* strains** | | |
| Af293 Δ*agd3* | Wild-type pathogenic strain of *A. fumigatus* with split marker, double homologous recombination to disrupt *agd3* | (12) |
| **Recombinant protein expression plasmids** | | |
| pET24a | IPTG-inducible expression vector encoding C-terminal hexahistidine tag, a thrombin cleavage site, and an optional C-terminal hexahistidine tag, Kan$^R$ | Novagen |
| pET28a | IPTG-inducible expression vector encoding N-terminal hexahistidine tag, a thrombin cleavage site, and an optional C-terminal hexahistidine tag, Kan$^R$ | Novagen |
| pET28a::*Pt*PelA$^{Δ36}$ | pET28a with *P. thermotolerans* H165_RS0111390 *pelA* corresponding to residues 37-937 fused to an N-terminal hexahistidine tag; Kan$^R$ | This study |
| pET28a::*Pt*PelA$^{D149A}$ | pET28a:: *Pt*PelA$^{Δ36}$ with a D149A mutation in the *pelA* gene | This study |
| pET28a::*Pt*PelA$^{E207A}$ | pET28a:: *Pt*PelA$^{Δ36}$ with a E207A mutation in the *pelA* gene | This study |
| pET28a::*Pt*PelA$^{D149A/E207A}$ | pET28a:: *Pt*PelA$^{Δ36}$ with D149A and E207A mutations in the *pelA* gene | This study |
| pET28a::*Pt*PelA$^{D517A}$ | pET28a:: *Pt*PelA$^{Δ36}$ with a D517A mutation in the *pelA* gene | This study |
| pET28a::*Pt*PelA$^{D519A}$ | pET28a:: *Pt*PelA$^{Δ36}$ with a D519A mutation in the *pelA* gene | This study |
| pET28a::*Pt*PelA$^{H589A}$ | pET28a:: *Pt*PelA$^{Δ36}$ with a H589A mutation in the *pelA* gene | This study |
| pET28a::*Pt*PelA$^{H593A}$ | pET28a:: *Pt*PelA$^{Δ36}$ with a H593A mutation in the *pelA* gene | This study |
| pET28a:: *Pa*PelA$^{Δ46}$ | pET28a with *P. aeruginosa* PAO1 *pelA* corresponding to residues 46-948 fused to an N-terminal hexahistidine tag; Kan$^R$ | (11, 13) |

| | | |
|---|---|---|
| pET28a::*Pa*PelA$^{D160A/E218A}$ | pET28a:: *Pa*PelA$^{\Delta46}$ with D160 and E218 mutations in the *pelA* gene | This study |
| pET28a::*Pa*PelA$^{hydr}$ | pET28a with *P. aeruginosa* PAO1 *pelA* corresponding to residues 47-303 fused to an N-terminal hexahistidine tag; Kan$^R$ | (14) |
| pET28a::*Pa*PelB$^{47\text{-}880}$ | pET28a with *P. aeruginosa* PAO1 *pelB* corresponding to residues 47-880 fused to an N-terminal hexahistidine tag; Kan$^R$ | (15) |

Kan, kanamycin.

**Table S7. Primers used in this study.**

| Name | Sequence (5' → 3') |
|---|---|
| **Recombinant protein purification** | |
| *Pt*PelA-*Pt*-37-F | GTC **CAT GGC** TAA ACC GTC TTC TGT TG |
| *Pt*PelA-*Pt*-937-R | GTG **GTG CTC** GAG GTT GCA AAC |
| *Pt*PelA-D149A-F | T CTG TTC CTG GcT ACT CTG GAC T |
| *Pt*PelA-D149A-R | A GTC CAG AGT AgC CAG GAA CAG A |
| *Pt*PelA-E207A-F | A GTT GCC GTT Gct AGC ATC CAC G |
| *Pt*PelA-E207A-R | C GTG GAT GCT agC AAC GGC AAC T |
| *Pt*PelA-D517A-F | G GTT CAC ATC GcT GGC GAC GGT T |
| *Pt*PelA-D517A-R | A ACC GTC GCC AgC GAT GTG AAC C |
| *Pt*PelA-D519A-F | C ATC GAT GGC Gct GGT TTC GTT A |
| *Pt*PelA-D519A-R | T AAC GAA ACC agC GCC ATC GAT G |
| *Pt*PelA-H589A-F | A GTT GCT TCT gct ACC TTC AGC C |
| *Pt*PelA-H589A-R | G GCT GAA GGT agc AGA AGC AAC T |
| *Pt*PelA-H593A-F | C ACC TTC AGC gct CCG TTC TTC T |
| *Pt*PelA-H593A-R | A GAA GAA CGG agc GCT GAA GGT G |
| **Sequencing** | |
| T7 | TAA TAC GAC TCA CTA TAG GG |
| T7ter | GCT AGT TAT TGC TCA GCG G |
| *PtpelA*-SEQ-int1 | GGT TAC GCA GGT CTG TTC C |
| *PtpelA*-SEQ-int2 | GGA TAC TCT GGG CGT TGG |
| *PtpelA*-SEQ-int3 | GAA TTC CAC GAT CAG AGC CTG |
| *PtpelA*-SEQ-int4 | GTA CCC ACT GCT GAC CTC |
| *PtpelA*-SEQ-int5 | GGC GGT AAC ACT ATG CTG AC |

*Restriction sites and Gateway att sequences are bolded; regions of complementary to the target amplicon are underlined; lowercase letters denote a nucleotide substitution; synthetic ribosomal binding sites are in bold italics.

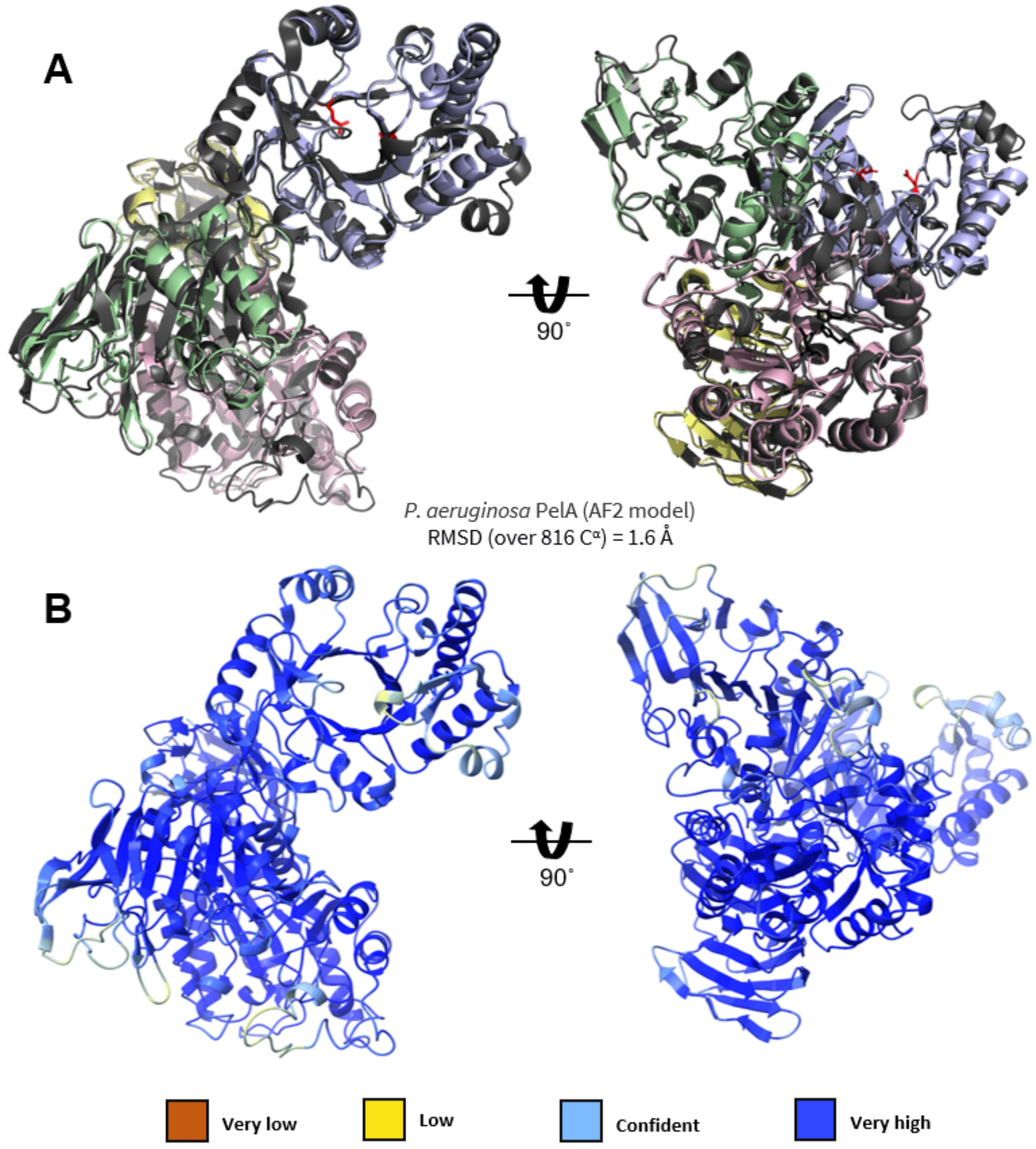


**Supplemental figure 1. The structure of *Pt*PelA aligns closely to the AF2 model of *Pa*PelA. (A)** Structural comparison of *Pt*PelA with the AF2 model of *Pa*PelA (grey). **(B)** Confidence levels of the AF2 prediction are mapped onto the *Pa*PelA model.

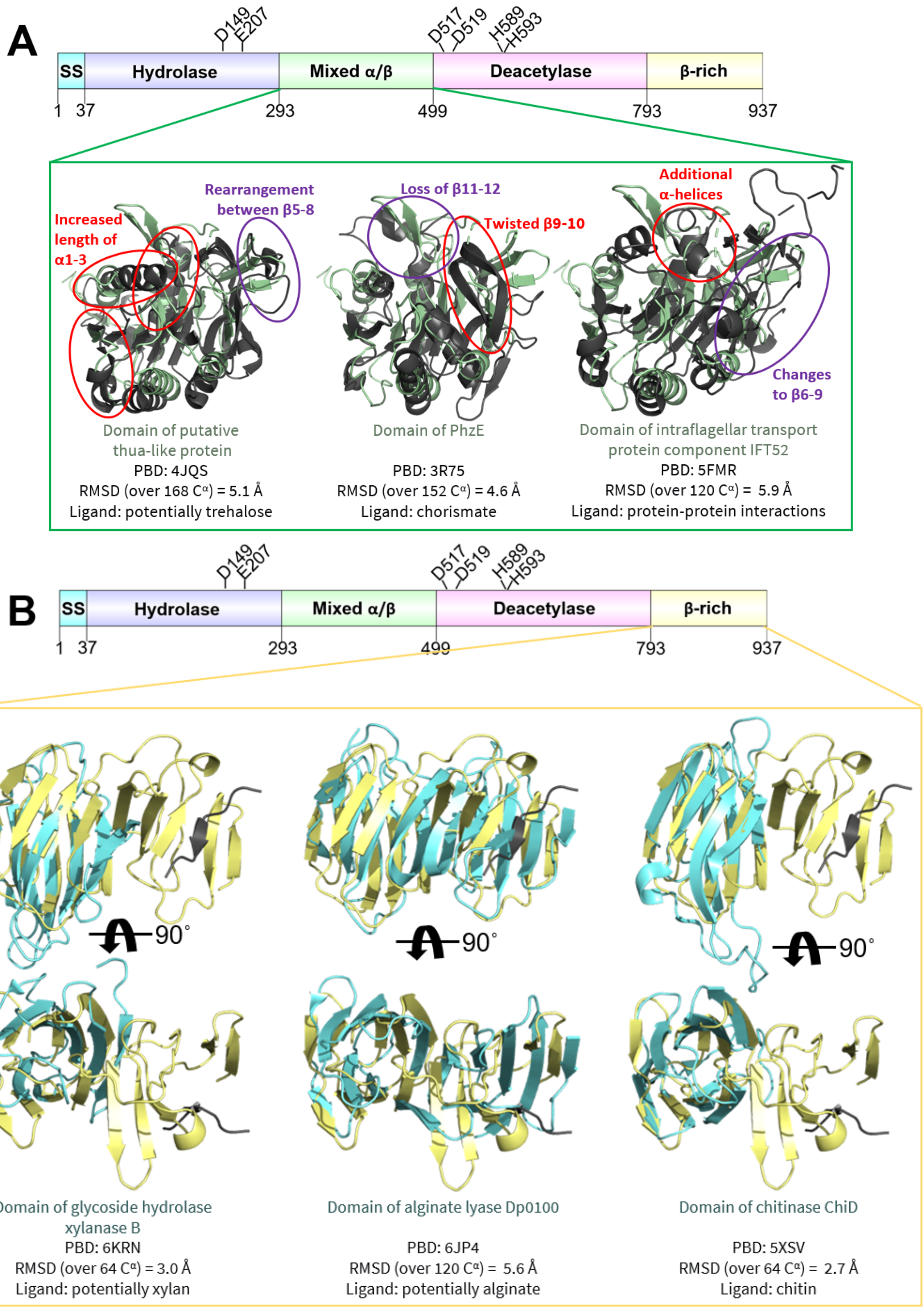


**Supplemental figure 2. Structural homologues of the mixed α/β and β-rich domains in *Pt*PelA have varied roles. (A)** Cartoon representation of the top three hits identified in a DALI search (dark grey) after superposition with the *Pt*PelA mixed α/β domain (green). **(B)** Superposition of the β-rich domain in *Pt*PelA (yellow) with three of the top hits identified in a DALI search (cyan) from two opposing views. The $C^{\alpha}$ atom RMSD values and the known or predicted ligands for the homologous structures are listed. SS, signal sequence.

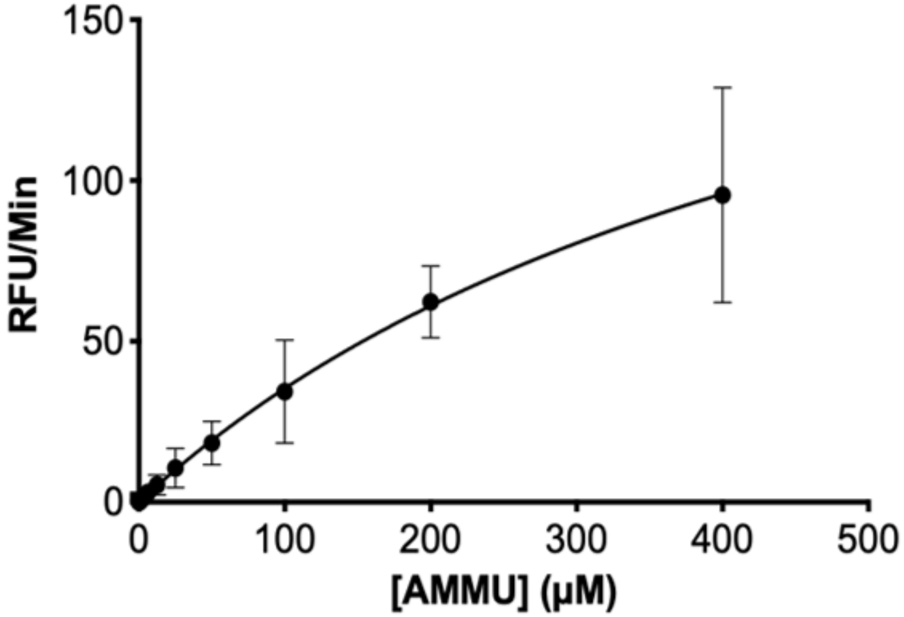


**Supplemental figure 3. *Pt*PelA has esterase activity.** Detection of *Pt*PelA esterase activity using AMMU as a pseudo-substrate. The error bars show the standard error of the mean for the three independent assays. AMMU, acetoxymethyl-4-methylumbelliferone.

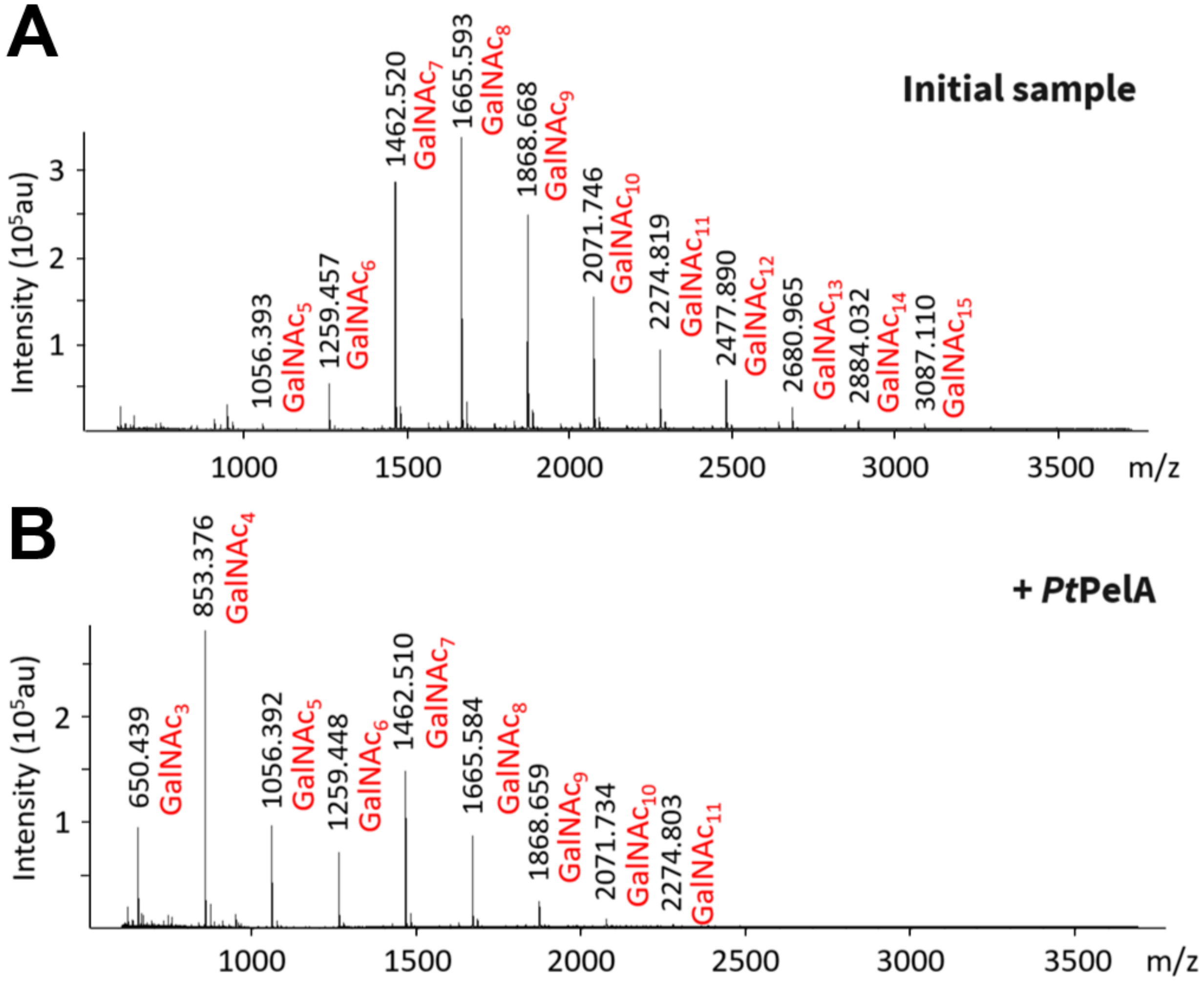


**Supplemental figure 4. *Pt*PelA has hydrolase activity. (A-B)** MALDI-TOF MS spectra of the initial pool of α-(1,4)-GalNAc oligosaccharides **(A)** and the products after incubation with *Pt*PelA **(B)** for 24 h.

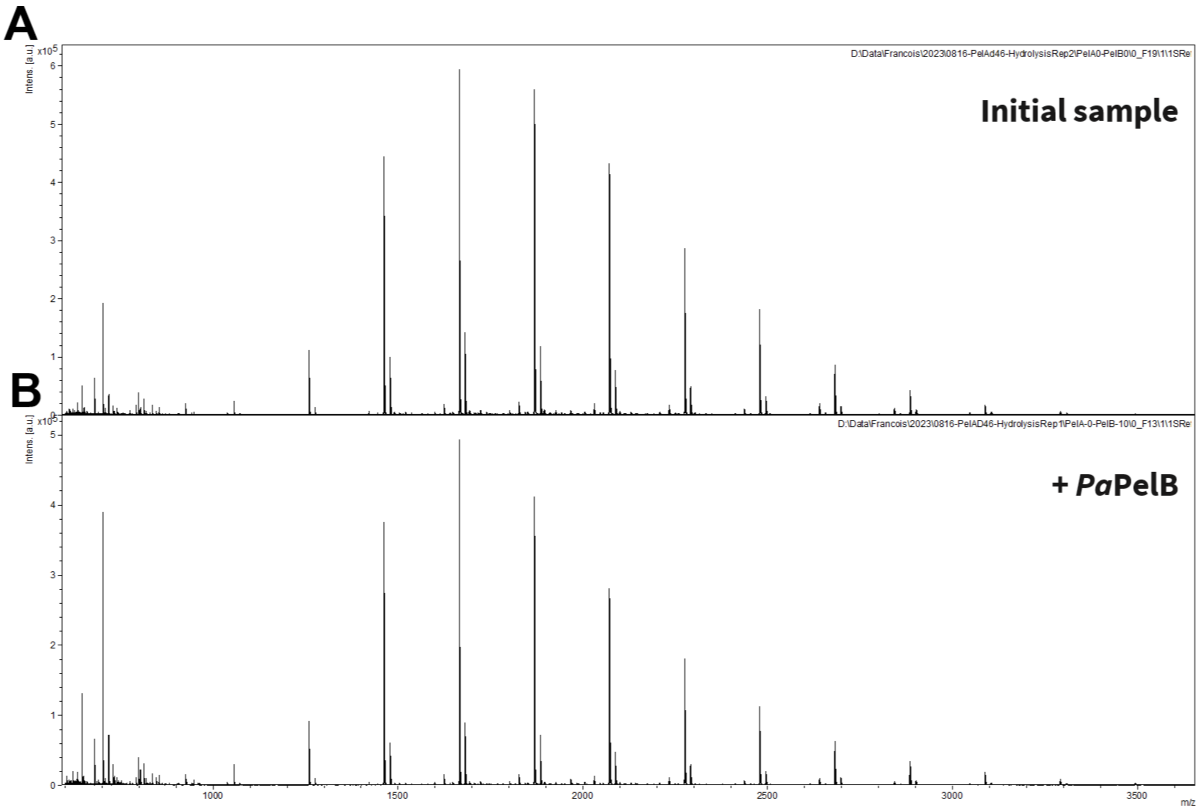


**Supplemental figure 5. *Pa*PelB does not modify α-(1,4)-GalNAc oligosaccharide. (A-B)** MALDI-TOF MS analysis of the initial pool of α-(1,4)-GalNAc oligosaccharides after incubation for 24 h in either the absence **(A)** or presence **(B)** of *Pa*PelB.

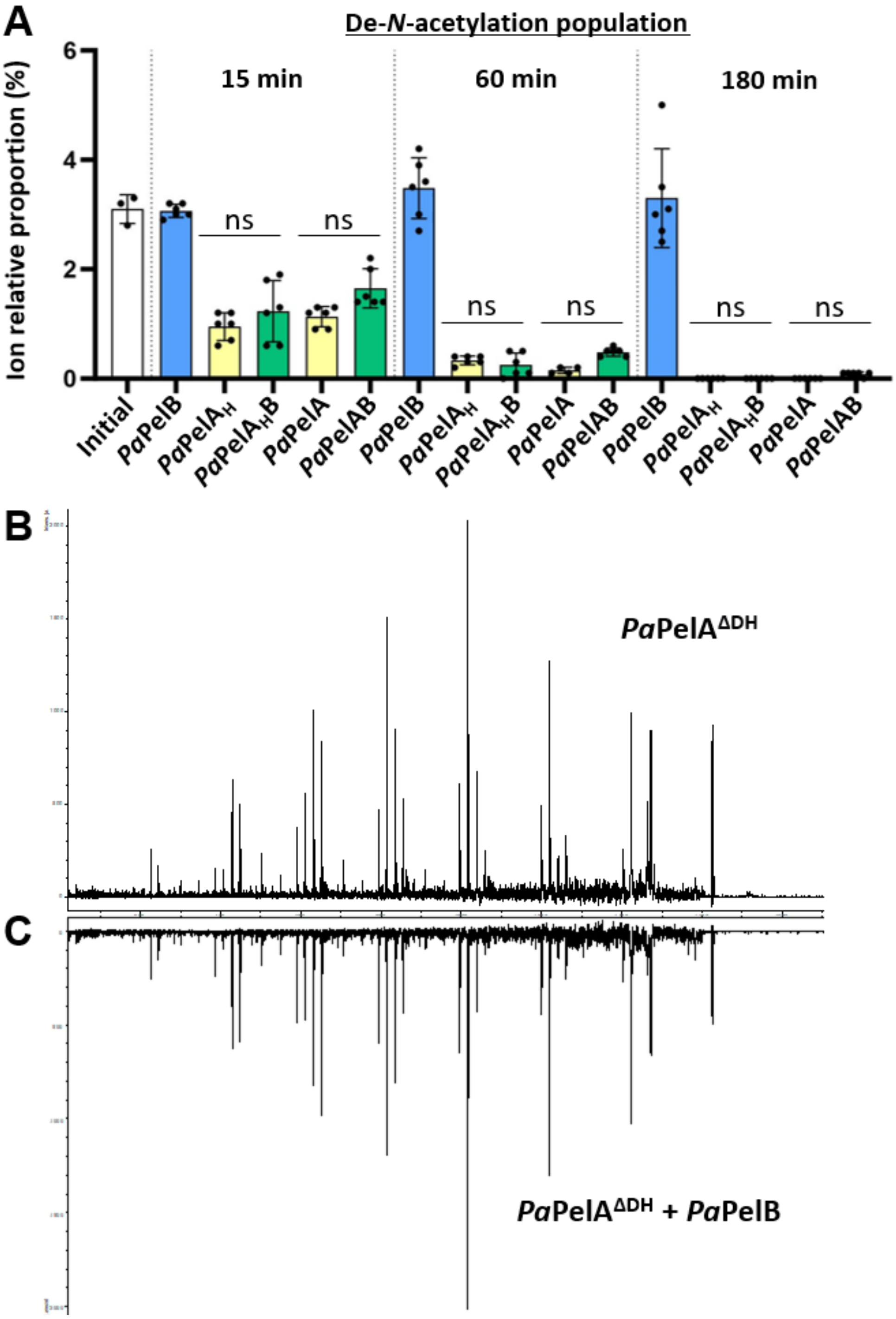


**Supplemental figure 6. Interaction of *Pa*PelB does not change the pattern of deacetylation by *Pa*PelA.** **(A)** The ion relative proportion of the MALDI-TOF MS enzyme spectra of the de-*N*-acetylation population after incubating wildtype *Pa*PelA or *Pa*PelA$_H$ in the absence or presence of *Pa*PelB. The data represent three biological replicates each with two technical replicates. Statistical significance was calculated using Kruskal Wallis multiple comparison tests between the indicated reaction conditions. Ns, not significant. **(B-C)** MS-MS spectra of an 8-mer oligomer with one de-*N*-acetylation produced from *Pa*PelA$^{\Delta DH}$ treatment in the absence **(B)** or presence **(C)** of *Pa*PelB.

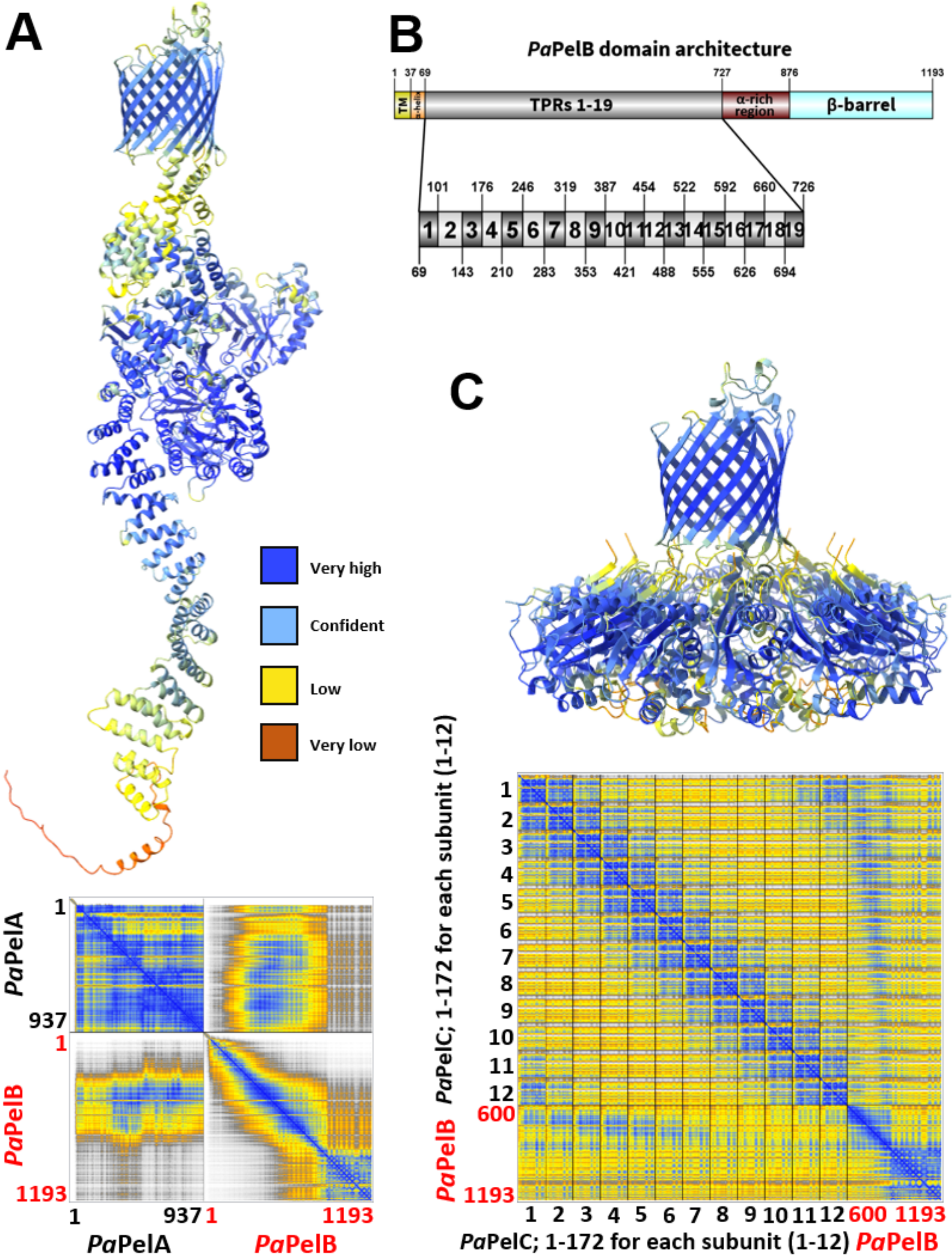


**Supplemental figure 7. Confidence score and PAE plot of the AF2 predicted *Pa*PelABC model. (A)** The AF2 structural prediction of the *Pa*PelAB complex from Fig. 8 is colored by the confidence level. The corresponding PAE plot is shown below. **(B)** The domain architecture of *Pa*PelB, as updated from the AF2 model prediction. TM, transmembrane; TPR, tetracopeptide repeats. **(C)** The AF2 structural prediction of the *Pa*PelBC complex from Fig. 8 is colored by the confidence level. The corresponding PAE plot is shown below.

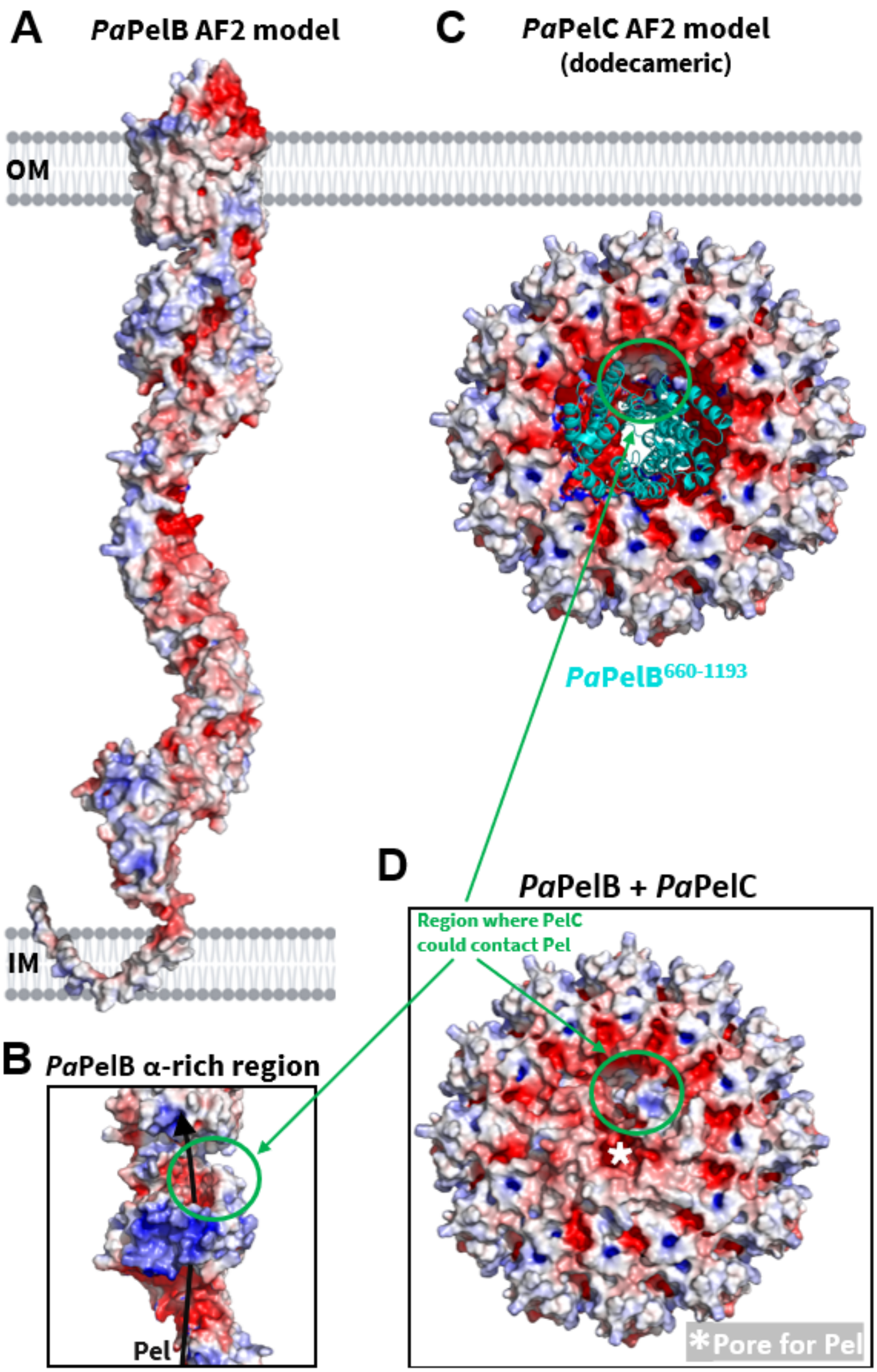


**Supplemental figure 8. AF2 structural models from Fig. 8 as colored by electrostatics. (A)** Electrostatic surface representation of *Pa*PelB calculated by APBS in PyMol and visualized from -5 (blue) to 5 (red) kT/e. **(B)** Electrostatic surface representation of the α-rich region of *Pa*PelB. **(C)** Electrostatic surface representation of dodecameric *Pa*PelC. Amino acids 660-1193 of *Pa*PelB from the *Pa*PelBC AF2 complex prediction are shown in cartoon representation (cyan). **(D)** Electrostatic surface representation of *Pa*PelBC. IM, inner membrane; OM, outer membrane.

### Supporting information methods

#### *Differential scanning fluorimetry (DSF)*

Purified wild-type and mutant *Pt*PelA were diluted to a concentration of 0.1 µM in 25 µL. These samples also included 5 µL of 1:500 SYPRO Orange. Samples were added to white conical-bottom 96-well plates (Bio-Rad) in triplicate. Samples were recorded from 25 to 80 ˚C on a real-time PCR machine (Bio-Rad CFX Connect) at a wavelength of 554 nm. Results were analyzed using Prism (GraphPad Software Inc.).

#### *AMMU esterase assay*

This assay was performed as previously described (16). Thawed wild-type and mutant *Pt*PelA was diluted to 1 µM in the assay buffer (50 mM HEPES pH 7 and 75 mM NaCl) and kept on ice. The protein was then diluted to 1.11 µM and 22.5 µL of protein was added to PCR strip tubes. The AMMU substrate was dissolved in DMSO and serially diluted from 4 mM to 4 µM. Reactions were initiated when 2.5 µL of AMMU was added to the enzyme reactions solutions to a total reaction volume of 25 µL. From the PCR strip tubes, 10 µL from the reaction solution were added to a 384-well black bottom plate (Corning 3820) in duplicate and centrifuged at 3000 RPM for 30 s. Reaction progress was monitored in real-time by measuring the RFU at 30-s intervals over 10 min at room temperature. The $\lambda_{emm}$ and $\lambda_{ext}$ used were 330 and 450 nm, respectively. The background hydrolysis was monitored and subtracted from the enzyme-catalyzed reactions. All assays were performed in quadruplicate using a BioTek Synergy Neo2 plate reader (Agilent Technologies). Prism (GraphPad Software Inc.) was used for all analyses.

**Supporting information references**
